# Live-cell RNA imaging with metabolically incorporated fluorescent nucleosides

**DOI:** 10.1101/2022.03.22.485351

**Authors:** Danyang Wang, Ana Shalamberidze, A. Emilia Arguello, Byron Purse, Ralph E. Kleiner

## Abstract

Fluorescence imaging is a powerful method for probing macromolecular dynamics in biological systems, however approaches for cellular RNA imaging are limited to the investigation of individual RNA constructs or bulk RNA labeling methods compatible primarily with fixed samples. Here, we develop a platform for fluorescence imaging of bulk RNA dynamics in living cells. We show that fluorescent bicyclic and tricyclic cytidine analogues can be metabolically incorporated into cellular RNA by overexpression of uridine-cytidine kinase 2 (UCK2). In particular, metabolic feeding with the tricyclic cytidine-derived nucleoside tC combined with quantitative confocal imaging enables the investigation of RNA synthesis, degradation, and trafficking at single-cell resolution. We apply our imaging modality to study RNA metabolism and localization during the oxidative stress response and find that bulk RNA turnover is greatly accelerated upon NaAsO_2_ treatment. Further, we identify cytoplasmic RNA granules containing RNA transcripts generated during oxidative stress that are distinct from canonical stress granules and P-bodies and co-localize with the RNA helicase DDX6. Taken together, our work provides a powerful approach for live-cell RNA imaging and reveals how cells reshape RNA transcriptome dynamics in response to oxidative stress.

## INTRODUCTION

RNA plays a central role in biology and probing its dynamic behavior is critical for illuminating the mechanisms underlying fundamental biological processes. A powerful approach to study the lifecycle of cellular RNA, including its transcription, trafficking, and decay is fluorescence microscopy(1); however live-cell RNA imaging remains a major challenge. Current approaches for visualizing RNA dynamics in living cells have primarily focused on imaging individual transcripts. This can be accomplished in several ways. Singer and co-workers developed MS2 tagging(2), which uses a viral MS2 stem-loop sequence to recruit a GFP-fusion protein to the RNA of interest, and has been the gold standard for live-cell RNA imaging over the last three decades. More recently, fluorescent RNA aptamers(3) and CRISPR-Cas targeting strategies(4) have provided complementary methods to image individual RNA sequences in cells. Further, RNAs generated *in vitro* or purified from cells can be chemoenzymatically modified with synthetic fluorophores and then introduced into cells(5). These methods for live-cell RNA imaging have provided important biological insights into the behavior of individual RNA transcripts, however, they lack the simplicity of fluorescent protein-fusions, and suffer from various drawbacks including low signal, high background fluorescence, and the necessity to introduce non-native RNA sequences into the transcript of interest. In addition, none of these methods can monitor global RNA dynamics in a living cell.

The metabolic incorporation of modified nucleosides into cellular RNA is a versatile approach to study RNA transcription and turnover(6). We and others have reported that modified pyrimidine and purine nucleosides containing alkyne(7–10), azide(11–14), or vinyl(15, 16) functionality can be incorporated into cellular RNA through nucleotide salvage pathways, labeled with fluorescent dyes using bioorthogonal chemistry, and used to visualize RNA in intact cells and organisms. Thus, unlike RNA imaging strategies relying upon exogenous sequence tags or recruitment of RNA-binding proteins, the power of metabolic labeling lies in its simplicity, ease of use, and transcriptome-wide generality. Azide/alkyne-modifications that minimally perturb nucleoside structure can be accepted by nucleotide salvage pathways and incorporated into RNA in living cells, however, imaging of these metabolically incorporated bioorthogonal ribonucleosides has been performed largely in fixed cells or tissue sections. This is primarily due to reliance on Cu(I)-catalyzed azide-alkyne cycloaddition (CuAAC)(17) for fluorescent labeling, which is incompatible with living cells as well as the need to “wash out” excess unreacted dye used to drive the labeling reaction to completion. While these approaches are powerful, bioorthogonal labeling protocols can introduce bias due to fixation and permeabilization methods, or as a result of non-specific binding of the fluorescent dye to cellular structures, and therefore it is unknown whether these methods accurately capture a snapshot of all cellular RNAs. In a few instances, strain-promoted azide-alkyne cycloaddition (SPAAC) chemistry has been utilized to label azido-modified RNA with fluorescent dyes in living cells(13, 14), however the slow kinetics of this reaction, requirement for cell permeable fluorescent dyes, and difficulty in removing excess fluorophore make challenging its application to intracellular targets.

An alternative to the two-step procedure used for imaging RNA with bioorthogonal ribonucleosides would be the direct metabolic incorporation of a fluorescent nucleoside (Fig. 1a). Since this strategy would not require a fluorophore labeling reaction after RNA incorporation and the fluorescence of an RNA transcript containing multiple modified nucleosides should exceed the fluorescence background from individual modified nucleosides or nucleotides, it should enable live-cell RNA imaging of global transcriptome dynamics. Indeed, a large number of fluorescent nucleoside analogs have been reported(18), however these structure have been primarily used for *in vitro* applications and incorporated into RNA using chemical synthesis or *in vitro* transcription, and it is unknown whether they are suitable substrates for metabolic incorporation into RNA.

**Figure 1.**
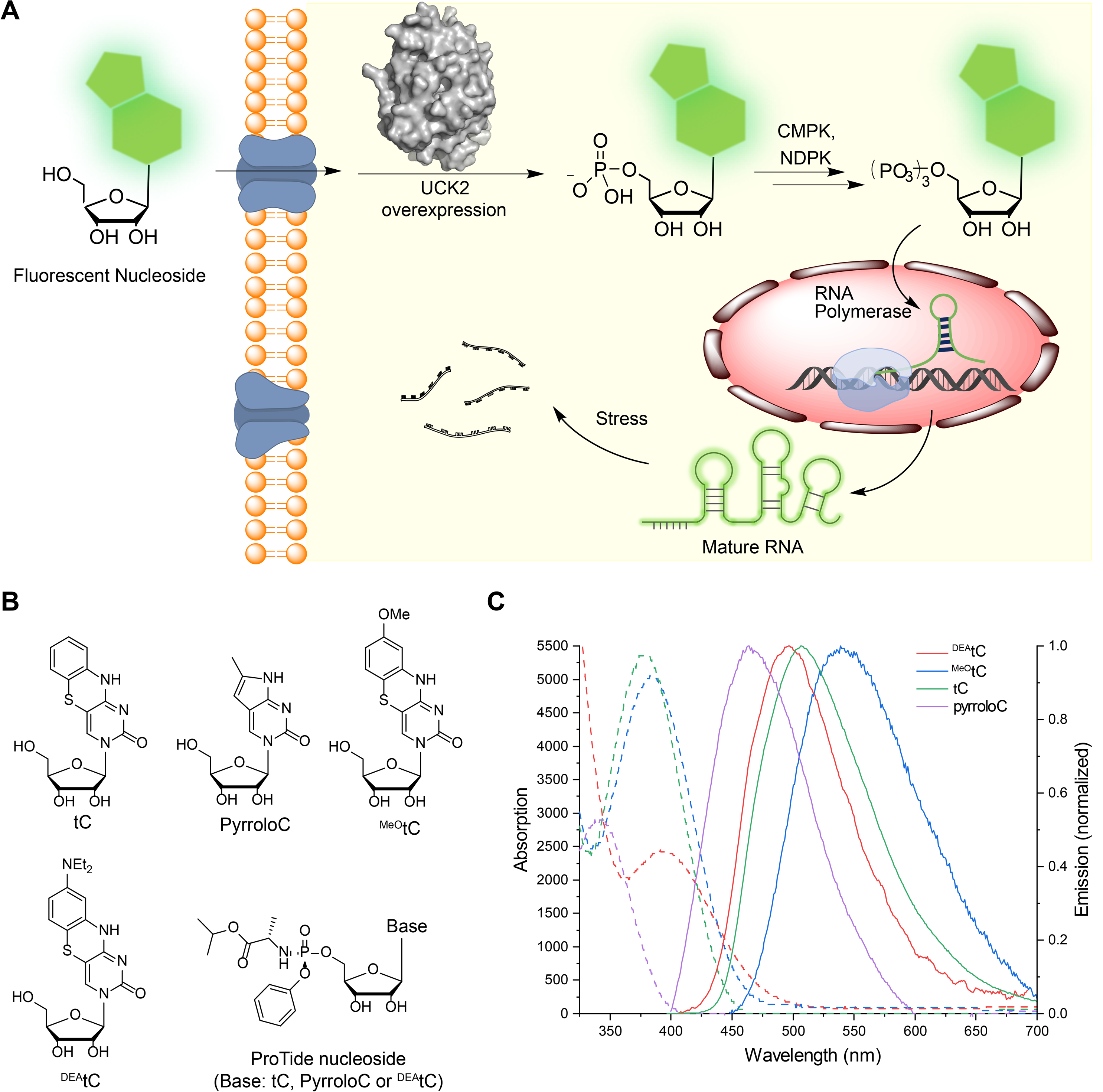
Metabolic incorporation of fluorescent nucleosides into cellular RNA. (A) Scheme for incorporation of fluorescent nucleosides through the nucleotide salvage pathway. (B) Structure of fluorescent nucleosides and ProTide nucleosides used in this work. (C) Absorption (dashed) and corrected emission (solid) spectra of fluorescent ribonucleoside analogues used in this study. Spectra were recorded at 295 K in 1× PBS buffer, pH 7.4.

Previously, we showed that phosphorylation by uridine-cytidine kinase 2 (UCK2) is the bottleneck for the metabolic incorporation of C5-modified pyrimidine nucleosides and that expression of a mutant form of the kinase enabled metabolic labeling with 5-azidomethyluridine (5-AmU)(13). Herein, we evaluate the *in vitro* phosphorylation and RNA metabolic incorporation of a series of fluorescent cytidine analogues. We demonstrate that expression of UCK2 in cells facilitates efficient RNA labeling with pyrrolocytidine (pyrroloC)(19) and 1,3-diaza-2-oxophenothiazine (tC)(20) fluorescent ribonucleosides. Further, we use the UCK2 system combined with tC feeding and confocal microscopy in order to measure RNA transcription, turnover, and trafficking in live cells during normal and stress conditions. We reveal the effects of oxidative stress on RNA metabolism and identify the formation of a novel stress-dependent RNA-protein granule. Taken together, our work provides a general approach for live-cell RNA imaging and reveals the spatiotemporal dynamics of cellular RNA during the oxidative stress response.

## RESULTS

### Fluorescent nucleosides for RNA imaging

In order to label cellular RNA for biological imaging, we selected and evaluated a set of fluorescent pyrimidine nucleosides. Our criteria included commercial availability or straightforward synthetic accessibility of the structure, ability of the modified nucleoside to engage in canonical Watson-Crick base pairing, likelihood of compatibility with pyrimidine salvage pathways, and fluorophore brightness and spectral overlap with commonly used imaging modalities in fluorescence microscopy. Therefore, we chose four modestly sized bicyclic and tricyclic fluorescent cytidine analogues containing extended ring systems projecting from the non-Watson Crick face of the molecule (Fig. 1b). We measured the absorption and emission spectra and determined the fluorescence quantum yield (*Φ*_em_) of these analogues under identical conditions – dilute solutions in 1× PBS buffer (pH 7.4) at 22 °C (Fig. 1c, Supplementary Table 1). PyrroloC(19, 21, 22) is a commercially available bicyclic cytidine analogue for which we measured ε_357 nm_ = 2900 M^−1^ cm^−1^ and *Φ*_em, 465 nm_ = 0.54. It has the smallest structural deviation from natural pyrimidine nucleosides among this set, but is significantly quenched by base pairing and stacking, with reported *Φ*_em, 460 nm_ = 0.06 in duplex DNA or RNA(23). The remaining three compounds are members of the tC family(20). The tC ribonucleoside was synthesized following published procedures(24), and we developed analogous methods to synthesize ^MeO^tC and ^DEA^tC ribonucleosides, which are reported here (compound synthesis and characterization data are available in the Supplementary Information). Parent tC offers robust fluorescence as a free nucleoside with ε_377nm_ = 5400 M^−1^ cm^−1^ and *Φ*_em, 513 nm_ = 0.17, and similar brightness in matched duplex DNA, *Φ*_em, 513nm_ = 0.11(20, 25, 26). ^DEA^tC has little fluorescence as a free nucleoside (*Φ*_em,493nm_ = 0.006) but much higher quantum yield up to *Φ*_em, 500 nm_ = 0.12 in dsDNA or greater in a DNA–RNA heteroduplex (up to *Φ*_em_ = 0.20; unpublished)(27–29). ^MeO^tC lands in the middle, modestly brighter than ^DEA^tC as a free nucleoside (*Φ*_em, 550 nm_ = 0.015) but with less fluorescence increase upon matched base pairing and stacking in dsDNA (*Φ*_em, 532 nm_ ≈ 0.03)(27). The insensitivity of parent tC fluorescence to local environment is attractive, while ^DEA^tC fluorescence turn-on upon incorporation into a duplex offers potential utility for probing nucleic acid structure in cells.

### Metabolic labeling with fluorescent cytidine analogues

We next evaluated whether the fluorescent nucleosides could be incorporated into living cells. For this purpose, we used HeLa cells stably expressing ribonucleoside kinase UCK2 under the control of a tetracycline-inducible promoter (i.e. Flp-In T-Rex system) or made to constitutively express UCK2 through transient transfection of a suitable DNA plasmid construct, and monitored incorporation using fluorescence microscopy and quantitative nucleoside LC-QQQ-MS. We reasoned that overexpression of WT or mutant UCK2(13) protein could be applied to facilitate the phosphorylation and incorporation of fluorescent nucleosides, in the case that these modified structures are incompatible with native nucleotide metabolism.

To test nucleoside incorporation, we treated cells with each fluorescent nucleoside for 12 hr and then imaged cells by epifluorescence microscopy after removing free nucleoside. PyrroloC, tC, and ^MeO^tC were used at 500 μM; ^DEA^-tC was used at 200 μM due to cytotoxicity. We chose a DAPI filter set (excitation: 360 ± 20 nm, emission: 460 ± 25 nm) due to its availability on most fluorescence microscopes and since the excitation and emission band pass wavelengths overlapped with the spectra of each of the synthetic nucleosides. Since we could not detect fluorescence in native HeLa cells after feeding (Supplementary Fig. 1), we tested whether induction of UCK2 expression (WT or Y65G mutant) in the HeLa T-Rex Flp-In UCK2 cells could facilitate nucleoside incorporation. Gratifyingly, in cells treated with tC or pyrroloC and induced to express WT or Y65G UCK2, we observed strong fluorescence labeling (Fig. 2a and Fig. 2b). Fluorescence signal was concentrated in the nucleus, but was also found in the cytoplasm, consistent with RNA incorporation. Interestingly, for both pyrroloC and tC, we observed higher fluorescence in cells overexpressing WT UCK2 as compared to the Y65G mutant, in contrast to our previous results with C5-modified uridine analogues containing linear substitutions. In cells without UCK2 overexpression, we detected minimal fluorescence (52-fold less) (Fig. 2a and Fig. 2b), indicating that endogenous UCK2 levels are insufficient for robust incorporation of these fluorescence nucleosides. For ^MeO^tC and ^DEA^tC derivatives, we were not able to observe fluorescent cellular labeling under any of the conditions tested. In addition, ^DEA^tC displayed significant cytotoxicity after extended treatment time.

**Figure 2.**
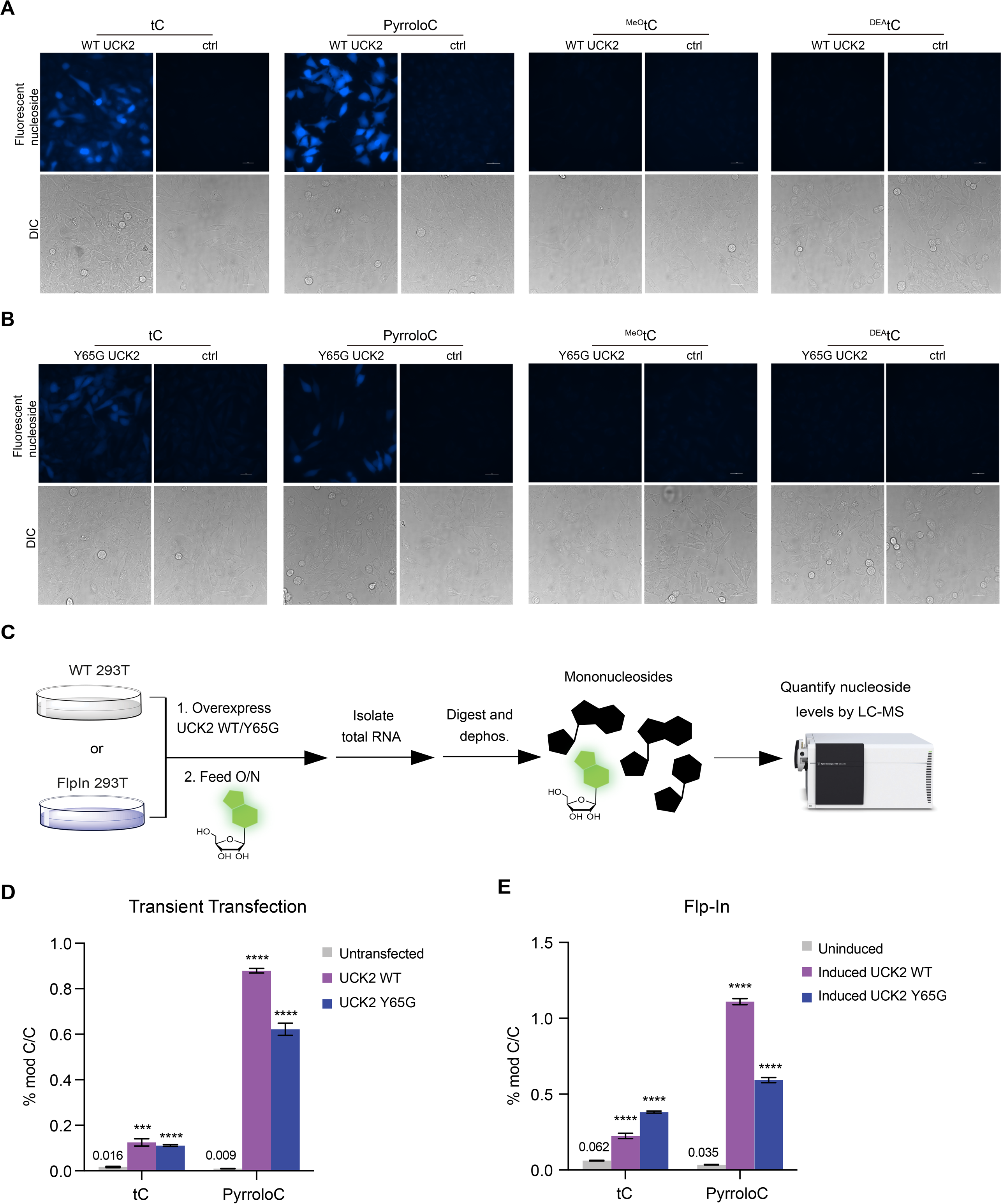
Detection of fluorescent nucleosides incorporation into cellular RNA using imaging and LC-QQQ-MS. (A) Incorporation of fluorescent nucleosides into HeLa Flp-In WT UCK2 cells. UCK2 expression was induced with tetracycline and cells were treated with 500 μM nucleoside for 12 h and imaged. Tetracycline induction was omitted to serve as a control for UCK2 expression. (B) Incorporation of fluorescent nucleosides into HeLa Flp-In Y65G UCK2 cells. Cells were treated and imaged as in (A). (C) Workflow for quantitation of fluorescent nucleosides in cellular RNA using LC-QQQ-MS. (D) RNA incorporation of tC or pyrroloC into HeLa cells transfected with UCK2 constructs. (E) RNA incorporation of tC or pyrroloC in HeLa cell lines stably expressing inducible UCK2 WT or Y65G mutant.

To prove that cellular fluorescence is a result of RNA incorporation of pyrroloC and tC, rather than simply nucleoside uptake and phosphorylation, we employed two complementary assays. First, we performed feeding in the presence of actinomycin D (ActD), a potent inhibitor of RNA polymerase I and II. Under these conditions, cellular fluorescence was greatly reduced (15-fold less) (Supplementary Fig. 2). In contrast, co-treatment with hydroxyurea (a DNA synthesis inhibitor) had little effect on labeling. In addition, we isolated total cellular RNA from induced HeLa T-Rex Flp-In UCK2 cells or cells transiently transfected with UCK2 and performed nucleoside LC-QQQ-MS after digestion and dephosphorylation to nucleosides (Fig. 2c and Supplementary Table 2). Strikingly, our results show that overexpression of WT or Y65G UCK2 kinase in either expression system leads to ~20-100-fold increase in RNA incorporation of pyrroloC (Fig. 2d, Fig. 2e, Supplementary Fig. 5, Supplementary Table 3 and Supplementary Table 4). The highest level of pyrroloC labeling (1.1 ± 0.02% of C) was achieved using cells stably expressing inducible WT UCK2 (Fig. 2e). Similarly, we observed a ~4-8-fold increase in RNA labeling with tC upon expression of WT UCK2 or Y65G UCK2, although the overall levels of incorporation were slightly lower than those achieved with pyrroloC nucleoside (Fig. 2d and Fig. 2e)). We also measured incorporation of ^DEA^tC and ^MeO^tC using our LC-QQQ-MS assay. We were unable to detect ^DEA^tC nucleoside in digested total RNA (Supplementary Fig. 3, Supplementary Fig. 4, and Supplementary Table 5), consistent with our cellular imaging experiments (Fig. 2a and Fig. 2b). While we could measure 0.026 ± 0.0009% incorporation of ^MeO^tC upon UCK2 induction (Supplementary Fig. 4 and Supplementary Table 5), this level is ~10-40-fold lower than with pyrroloC or tC, and likely insufficient to produce robust fluorescence in cells. Taken together, our data demonstrates that efficient metabolic RNA labeling with fluorescent nucleosides pyrroloC and tC can be achieved through overexpression of nucleoside kinase UCK2. This system should be suitable for live-cell imaging of the bulk RNA population.

### In vitro phosphorylation with UCK2

To further understand the incorporation of bulky fluorescent nucleosides into cellular RNA and the involvement of UCK2 in this process, we directly studied the phosphorylation of modified nucleosides by recombinant, purified UCK2 using an HPLC-based *in vitro* assay. As we observed similar RNA incorporation levels in cells expressing either WT or Y65G UCK2, we focused on WT UCK2 since its heterologous expression and purification is more straightforward. Consistent with high-level cellular incorporation, we observed efficient phosphorylation of pyrroloC and tC nucleosides by UCK2 with complete phosphorylation of pyrroloC and 42% phosphorylation of tC in 1 hour under the conditions assayed (Fig. 3a, Fig. 3b, Supplementary Fig. 6, and Supplementary Table 6). Similarly, cytidine analogues that were poorly incorporated showed minimal *in vitro* phosphorylation -- we detected no phosphorylation of ^DEA^tC (even after extended reaction times) and only minimal phosphorylation (<20%) of ^MeO^tC after 1 hr (Fig. 3c, Fig. 3d, and Supplementary Table 6). Our results show that the incorporation efficiency of bicyclic and tricylic fluorescent cytidine analogues into cellular RNA correlates strongly with their ability to be phosphorylated by UCK2. Nevertheless, we cannot exclude that other factors including, but not limited to, nucleoside uptake, downstream metabolism, and toxicity of these nucleosides and related metabolites are also strong determinants of metabolic incorporation into cellular RNA.

In our previous work(13), we found that modest substitutions at the C5 position severely compromised the ability of uridine nucleosides to serve as substrates for UCK2, likely due to steric clash with aromatic residues in the catalytic center of the enzyme (e.g. Y65). Therefore, we were surprised that pyrroloC and tC were efficient substrates for UCK2 due to their large polycyclic structures. Since we had previously studied only C5-modified uridine nucleosides, we compared our results with pyrroloC and tC against two cytidine analogues containing different size substitutions at the C5 position: 5-bromocytidine (5-BrCyd) and 5-iodocytidine (5-ICyd). As expected, UCK2 activity on these substrates was inversely related to the size of the C5 substituent. Surprisingly, pyrroloC and tC were both more efficient UCK2 substrates than 5- BrCyd (Fig. 3e, Supplementary Fig. 7, and Supplementary Table 6), despite their multicyclic structure. We hypothesize that the rigidity of the pyrroloC and tC structures presents less steric bulk in proximity to UCK2 Y65, and indeed docking studies show that both nucleosides are well accommodated in the UCK2 active site (Fig. 3f and Supplementary Fig. 8). In addition, the extended π-systems of pyrroloC and tC could engage in productive stacking or Van der Waals interactions with aromatic residues in the UCK2 active site, such as Phe83 (Fig. 3f).

**Figure 3.**
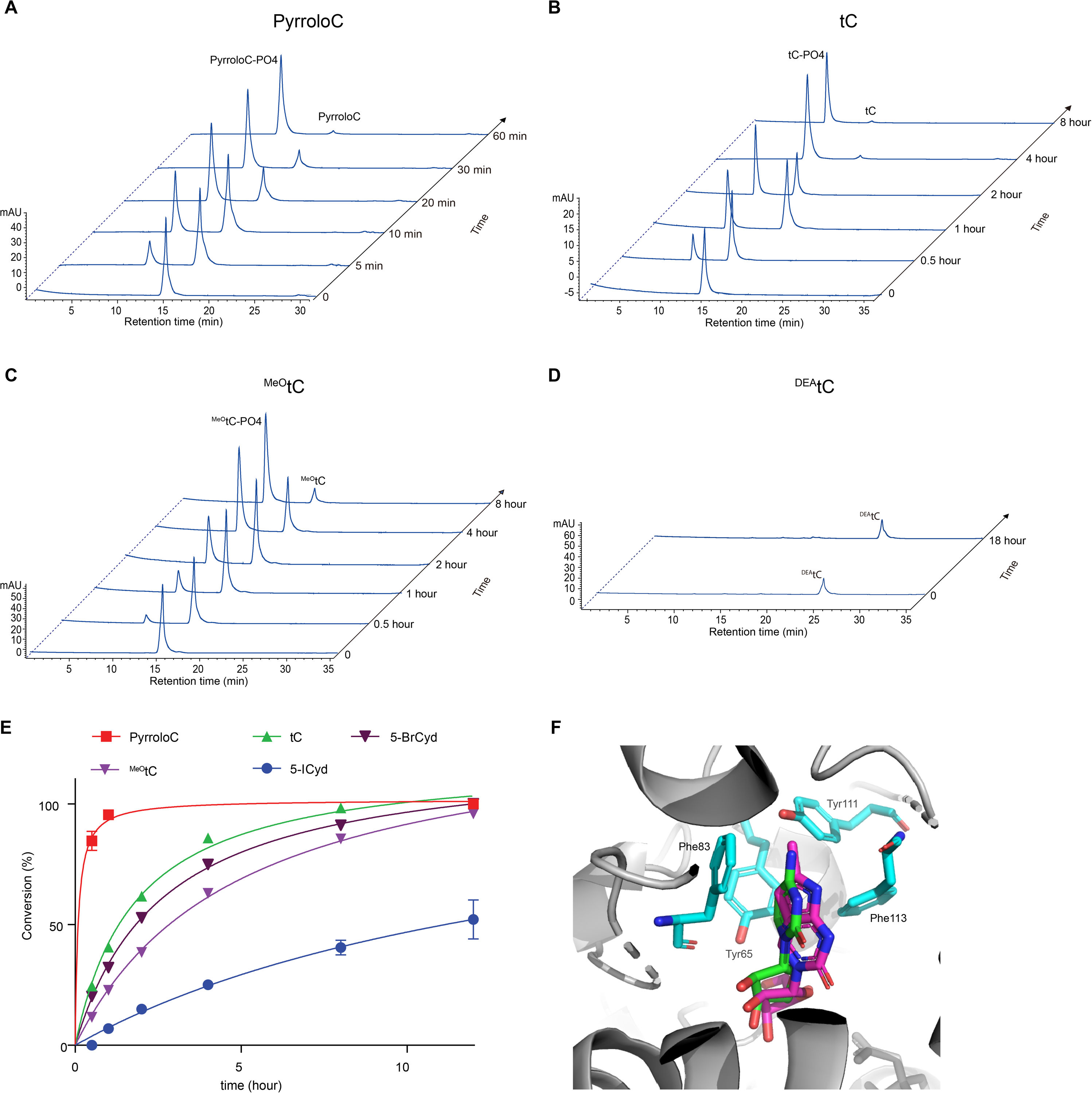
Phosphorylation of fluorescent nucleosides and cytidine analogues by recombinant UCK2. (A-D) HPLC chromatogram time course of PyrroloC (A), tC (B), ^MeO^tC (C) and ^DEA^tC(D) phosphorylation by recombinant UCK2 WT. (E) Quantification and kinetic analysis of fluorescent nucleoside phosphorylation reactions. Data represent mean ± SD (n = 3). (F). Overlay of simulated pyrroloC (purple) and cytidine (green) bound to UCK2 (PDB:1UEJ).

### ProTide fluorescent nucleoside derivatives are poorly incorporated into RNA

Due to the reliance of fluorescent nucleoside incorporation on UCK2 activity, we also sought to test a widely used kinase bypass method as a strategy to facilitate metabolic labeling. Kinase bypass approaches(30), which typically involve masked phosphate groups, allow nucleotides to be delivered across the plasma membrane and rely on subsequent enzymatic unmasking in the cytoplasm. One of the most successful of these approaches is the ProTide technology pioneered by McGuigan and used clinically in drugs including sofosbuvir and tenefovir alafenamide(31). We reasoned that ProTide versions of the fluorescent nucleosides might provide an alternative approach to incorporate these structures into nascent RNA without the need for UCK2 overexpression. Therefore, we prepared ProTide derivatives of pyrroloC, tC, and ^DEA^tC (Fig. 1b) and investigated their uptake into cells and incorporation into cellular RNA using fluorescence microscopy and LC-QQQ-MS. Of the three ProTide compounds assayed, we could detect cellular fluorescence only in cells treated with ProTide-tC (Supplementary Fig. 9a), suggesting that ProTide-pyrroloC and ProTide-^DEA^tC failed to penetrate cells, or were rapidly washed out before imaging. Interestingly, fluorescence resulting from ProTide-tC treatment was restricted to the cytoplasm and exhibited punctate staining (Supplementary Fig. 9a), which is inconsistent with RNA labeling. To further confirm that ProTide-tC was not incorporated into RNA, we co-treated cells with ActD and found no reduction in fluorescence (Supplementary Fig. 9b). Similarly, LC-QQQ-MS of isolated RNA from ProTide-tC treated cells indicated minimal incorporation – 51-fold lower than that achieved by treating UCK2-expressing cells with tC nucleoside (Supplementary Fig. 9c). While it does appear that ProTide-tC can diffuse across the plasma membrane, it is likely that unmasking is inefficient in HeLa cells, and further the compound appears to concentrate in cellular vesicles, precluding the use of the ProTide strategy for RNA metabolic labeling using tC and pyrroloC-derived nucleosides in this system. It is known from extensive work in medicinal chemistry that the effectiveness of phosphate group masking strategies is cell type-dependent(31, 32), and accordingly such fluorescent nucleotides may have utility in other cell types.

### Live cell RNA imaging

Efficient cellular RNA labeling with tC and pyrroloC upon UCK2 expression led us to pursue this system for imaging real-time RNA dynamics in living cells. Towards this goal, we treated UCK2-expressing HeLa cells with pyrroloC or tC and used confocal fluorescence microscopy to detect labeled RNA. First, we explored compatibility with different excitation sources and filter sets. For confocal imaging, 405 nm excitation combined with a DAPI emission filter gives similar signal intensity for both pyrroloC and tC; however, tC can also be imaged using 405 nm excitation combined with a GFP emission filter (535 ± 20 nm) due to its longer wavelength emissions, and indeed this combination gives better signal and allowed us to reduce exposure time in half (Supplementary Fig. 10a-c). Therefore, we decided to use the tC/UCK2 pair for confocal imaging with 405 nm excitation combined with GFP emission filter. First, we evaluated RNA labeling with different serum concentrations and in the presence of the RNA synthesis inhibitor actinomycin D (ActD). We treated cells with 200 μM tC, a concentration that balances cellular toxicity with incorporation efficiency (Supplementary Fig. 11), and monitored fluorescence after 6 hr. We observed tC fluorescence in the cytoplasm and nucleus with enhanced signal at the nucleolus, the major site of rRNA synthesis (Supplementary Fig. 10d), consistent with incorporation into RNA transcripts. Fluorescence was decreased substantially with ActD co-treatment (Fig. 4a), indicating that the majority of signal originates from tC-containing RNA transcripts as opposed to fluorescent tC metabolites (nucleotides). In addition, we found a 59% increase in fluorescence signal when tC treatment was performed in the presence of 10% fetal bovine serum as compared to reduced serum concentrations (Fig. 4a and Fig. 4b), in line with reports that RNA transcription is upregulated with increased growth rate(33). Next, we investigated the kinetics of RNA labeling with tC. We observed cellular fluorescence at time points as early as 30 min (Fig. 4c and Fig. 4d), indicating that tC RNA labeling can be used to report on dynamic cellular processes. Fluorescence signal increased over time and reached a plateau at 6 hr (Fig. 4c and Supplementary Fig. 10d).

**Figure 4.**
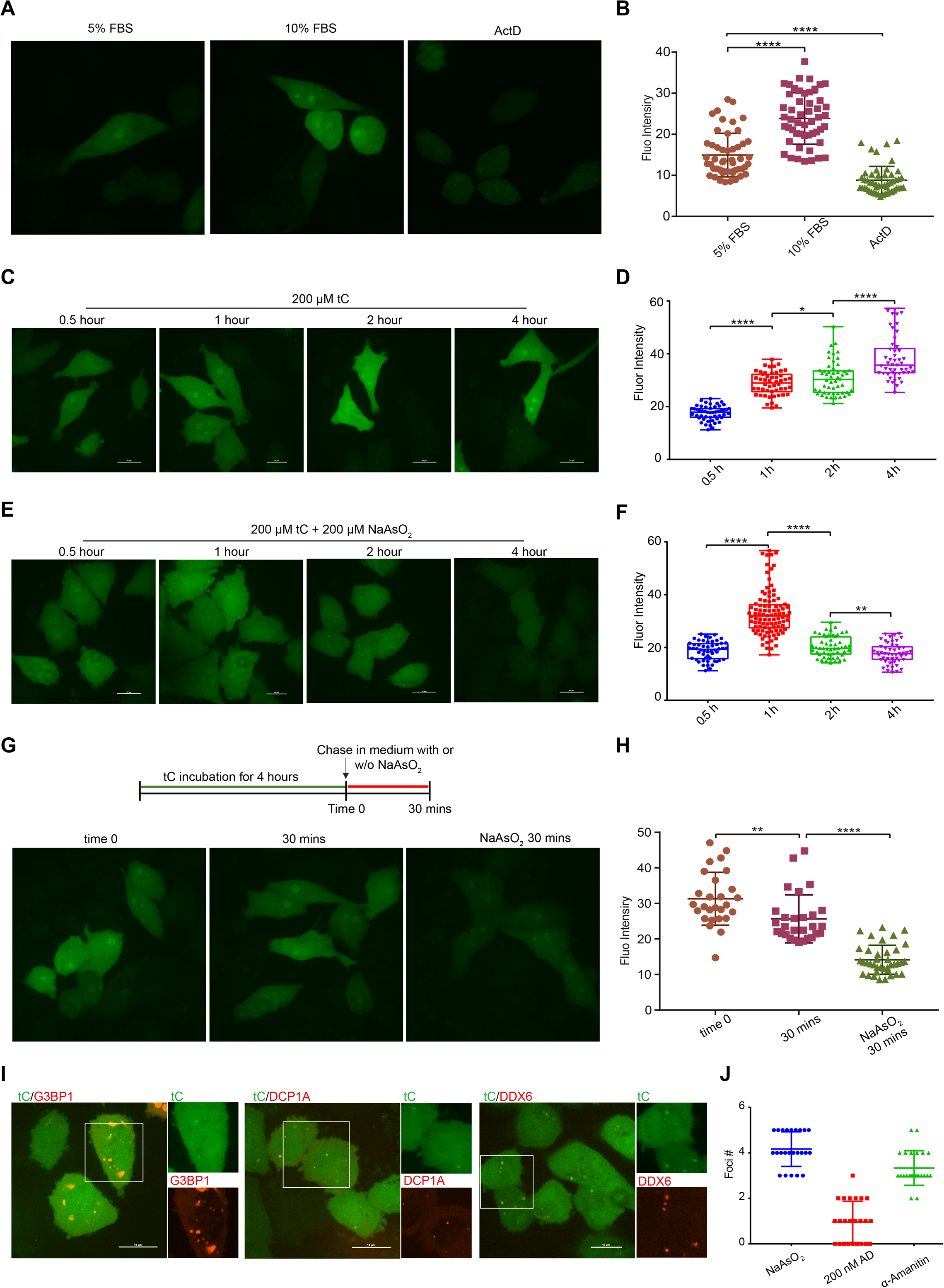
Live cell imaging of cellular RNA dynamics using tC RNA labeling. (A) Representative images of tC incorporation in cells grown in 5% or 10% FBS or in the presence of ActD. (B) Fluorescence intensity quantification of cells treated in (A). Error bars represent mean□±□s.d (n=□50 cells from three independent biological replicates). (C) Time-course analysis of tC incorporation into cellular RNA. (D) Fluorescence intensity quantification of cells treated in (C). Error bars represent mean□±□s.d. (n=□50 cells from three independent biological replicates). (E) Time-course analysis of tC incorporation into cellular RNA during co-treatment with 200 μM NaAsO_2_. (F) Fluorescence intensity quantification of cells treated in (E). Error bars represent meanL±Ls.d. (n=L50 cells from three independent biological replicates). (G) Representative images of cells pulsed with tC for 4 h and chased with complete medium or with 200 μM NaAsO_2_ in medium. (H) Fluorescence quantification of cells treated in (G). Error bars represent mean□±□s.d. (n=□25 cells from three independent biological replicates). (I) Colocalization of RNA foci formed under stress conditions with G3BP1, DCP1A or DDX6. (J) Quantification of cellular RNA foci number under stress and RNA synthesis inhibitors treatments. Error bars represent meanL±Ls.d. (n=L25 cells from three independent biological replicates).**P < 0.01, ****P < 0.0001.

Next, we used tC RNA labeling to investigate global RNA synthesis and turnover during oxidative stress. Oxidative stress induces a number of changes in cellular physiology concomitant with alterations in translational and transcriptional programs(34, 35), but the impact of oxidation on bulk RNA dynamics is still poorly understand. To investigate RNA dynamics during stress conditions we performed metabolic labeling with tC nucleoside during sodium arsenite-induced oxidative stress and imaged cells at multiple time points after initiating feeding and stress. We observed a biphasic response during arsenite stress – tC fluorescence increased during the first hr of treatment but then steadily decreased from 1 hr to 4 hr of continuous arsenite exposure (Fig. 4e and Fig. 4f). In contrast, tC accumulation in RNA in unstressed cells continuously increases during the course of 4 hr (Fig. 4c and Fig. 4d). In addition, NaAsO_2_-treated cells displayed changes in nucleolar RNA signal, exhibiting larger numbers of nucleoli with reduced size and more regular spherical shape as compared to irregularly shaped nucleoli in unstressed cells (Fig. 4e). Since NaAsO_2_ exposure could affect both RNA synthesis and degradation, we next performed a pulse-chase labeling experiment in order to specifically investigate RNA turnover. Cells were fed tC for 4 hr followed by a 30 min chase in tC-free medium in the presence or absence of NaAsO_2_. Fluorescence signal from tC-labeled RNA was strongly reduced (55%) during chase in AsO_2_-containing medium whereas tC-labeled RNA levels were only 18% reduced in unstressed cells during the 30 min chase (Fig. 4g and 4h). Taken together, our studies show that bulk RNA degradation rapidly increases in response to NaAsO_2_ stress, but they do not support an acute effect of arsenite stress on RNA synthesis. In addition, we find altered nucleolar RNA localization, which is suggests that oxidative stress affects rRNA synthesis and/or processing.

Interestingly, further analysis of RNA localization upon NaAsO_2_ treatment indicated the formation of cytoplasmic RNA foci that arose as early as 1 hr after stress induction (Fig. 4e, Fig. 4i and Supplementary Fig. 12). These foci were only present when NaAsO_2_ stress was applied during tC feeding, and were absent from unstressed cells (Fig. 4c) or cells where NaAsO_2_ was applied during the chase (i.e. after tC treatment) (Fig. 4g). To categorize the observed foci, we compared them against known cytoplasmic RNA-based structures specific to oxidative stress conditions. Stress granules are RNA-protein condensates containing translationally stalled mRNA and RNA binding proteins that form rapidly after arsenite stress(36). We tested whether our foci colocalized with the characteristic stress-granule protein G3BP1 using live-cell imaging and expression of G3BP1-mCherry together with tC labeling and NaAsO_2_ treatment. As expected, G3BP1-mCherry exhibited diffuse cytosolic staining during normal growth conditions and rapidly formed micron-sized foci in the cytosol upon NaAsO_2_ stress (Supplementary Fig. 13), but these foci did not co-localize with the tC-labeled RNA foci (Fig. 4i), although they were frequently found in close proximity. Next, we investigated co-localization with the P-body protein DCP1A(37). While P-bodies are present constitutively, their number has been shown to increase upon NaAsO_2_ stress(38). We imaged DCP1A-mCherry together with tC-labeled RNA, but did not observe co-localization between the NaAsO_2_-dependent RNA foci and P-bodies (Fig. 4i and Supplementary Fig. 13). Finally, we combined tC RNA live cell imaging with expression of the RNA helicase DDX6(39) and found that tC RNA foci consistently co-localized with DDX6-mCherry upon NaAsO_2_ stress (Fig. 4i and Supplementary Fig. 13). Further, to investigate the nature of the RNA in these foci, we tested their sensitivity to different small-molecule RNA polymerase inhibitors. Foci formation was unaffected by α-amanitin, an RNA polymerase II inhibitor, but abolished with ActD treatment at concentrations specific for RNA polymerase I inhibition (Supplementary Fig. 12). Taken together, our results show that a subset of RNA transcripts generated during arsenite stress accumulate in cytoplasmic foci together with DDX6. These foci are distinct from canonical stress granules and P-bodies and likely contain rRNA.

## DISCUSSION

In this manuscript we develop a strategy for whole transcriptome RNA imaging in living cells. We show through quantitative confocal fluorescence microscopy and nucleoside LC-QQQ-MS that overexpression of UCK2 enables metabolic RNA labeling and direct fluorescence imaging with fluorescent ribonucleoside cytidine analogs tC and pyrroloC. Compared to prior approaches that typically rely upon metabolic labeling with azide- or alkyne-modified nucleosides followed by bioorthogonal conjugation with a suitable reactive fluorophore, our strategy is operationally simpler, compatible with live-cell imaging, and less prone to artifacts resulting from cellular fixation and/or permeabilization and washing conditions.

We apply tC RNA labeling and confocal imaging in order to study the dynamics of RNA synthesis and turnover during the oxidative stress response. Our results show a dramatic and rapid increase in RNA turnover during sodium arsenite treatment, with a ~50% reduction in bulk RNA transcript levels transcribed before stress induction after only 30 minutes. In contrast, RNA synthesis rates appear largely unchanged in this initial period but then decline after more prolonged exposure to sodium arsenite. These RNA-specific effects likely play an important role in reshaping gene expression programs during the integrated stress response(34), both by downregulating the abundance of individual mRNA transcripts and by affecting rRNA levels and ribosome availability. In addition, we identified the formation of cytoplasmic arsenite-induced RNA granules that colocalize with the RNA helicase DDX6 but do not colocalize with the stress granule marker G3BP1 or the P-body protein DCP1A. Since these granules were only formed when tC RNA labeling was performed in the presence of NaAsO_2_ and not during a NaAsO_2_ chase, we propose that they contain RNA transcripts that were synthesized during stress conditions. Further, we find that granule formation is sensitive to the RNA Pol I inhibitor ActD but largely insensitive to the RNA Pol II inhibitor α-amanitin, suggesting that granules contain rRNA. Interestingly, a recent study demonstrated defects in 45S rRNA processing upon sodium arsenite exposure(40). We speculate that the accumulation of rRNA in cytoplasmic granules distinct from stress granules and P-bodies may be related to the elimination of misprocessed forms generated during oxidative stress. The accumulation of DDX6 in these structures is intriguing as DDX6 has been implicated in P-body assembly, mRNA decay, and translational repression(37, 41–43), but to our knowledge is not known to interact with rRNA. Further characterization of these granular structures will be necessary to understand their components and formulate hypothesis relating to their physiological relevance.

Our work demonstrates that fluorescent nucleosides can be incorporated into RNA through metabolic labeling and used to track RNA metabolism and intracellular trafficking. The key to this technology is judicious selection of nucleoside structure and overexpression of the ribonucleoside kinase UCK2. The bicyclic and tricyclic cytidine analogs explored herein likely represent a privileged scaffold for RNA metabolic labeling, and further investigation of this class of modified nucleosides together with UCK2 engineering should lead to the development of fluorescent nucleosides with enhanced and varied spectral properties and context-dependent fluorescence. Esbjorner and Wilhelmsson recently investigated the biological properties of synthetic mRNA containing the related tC° fluorescent nucleobase analogue(44), and found little effect on protein translation, underscoring the compatibility of these structures with biological systems. Beyond tracking localization and metabolism, analogous approaches could be utilized to incorporate diversely functionalized nucleoside analogs as biophysical probes to explore nucleic acid structure and protein-RNA interactions in their native context. Indeed, nucleic acid chemists have reported a variety of fluorescent nucleobases(18), but to our knowledge these structure have been primarily utilized in the context of oligonucleotides generated synthetically or by *in vitro* transcription. We envision that integrating nucleoside chemistry, protein engineering, and metabolic labeling should lead to unique opportunities for probing the dynamic behavior of RNA in its native biological context.

## ACKNOWLEDGEMENTS

We thank Gary Laevsky at the Princeton University Confocal Imaging Facility for assistance with live-cell fluorescence microscopy. We are grateful to David Sanders and Clifford Brangwynne for providing expression plasmids for G3BP1, DDX6, and DCP1A. R.E.K. acknowledges support from a National Science Foundation CAREER award (MCB-1942565), the National Institute of Health (R01 GM132189), and the Alfred P. Sloan Foundation. B.W.P. acknowledges the National Science Foundation (CHE-1800529 and CHE-2102642) for financial support. A.E.A. was supported by an Eli Lilly-Edward C. Taylor Fellowship in Chemistry.

## AUTHOR CONTRIBUTIONS

R.E.K. and B.W.P. conceived the idea and directed research. D.W. performed biochemical experiments and cellular imaging experiments. A.S. synthesized fluorescent nucleosides and fluorescent ProTide compounds. A.E.A. performed nucleoside LC-QQQ-MS experiments. All authors contributed to the preparation of the manuscript.

