## Supplementary Information for "Live-cell RNA imaging with metabolically incorporated fluorescent nucleosides"

Table of Contents


i. Methods
ii. Supplementary Table 1
iii. Supplementary Table 2
iv. Supplementary Table 3
v. Supplementary Table 4
vi. Supplementary Table 5
vii. Supplementary Table 6
viii. Supplementary Figure 1
ix. Supplementary Figure 2
x. Supplementary Figure 3
xi. Supplementary Figure 4
xii. Supplementary Figure 5
xiii. Supplementary Figure 6
xiv. Supplementary Figure 7
xv. Supplementary Figure 8
xvi. Supplementary Figure 9
xvii. Supplementary Figure 10
xviii. Supplementary Figure 11
xix. Supplementary Figure 12
xx. Supplementary Figure 13
xxi. Chemical synthesis
xxii. References
xxiii. Appendix

### Experimental Methods

#### Chemicals and Synthesis

Starting materials and reagents were synthesized using reported procedures ($^{DEA}$tC nucleobase, tC nucleobase, $^{MeO}$tC nucleobase)[1] or purchased at ACS reagent grade or higher from Acros Organic, Sigma-Aldrich, Combi-Blocks, and Berry & Associates, and used without further purification. 5-Iodocytidine (5-IC) and 5-Bromocytidine (5-BrC) were purchased from Carbosynth. Analytical thin-layer chromatography was performed using pre-coated MilliporeSigma™ 200 μm silica gel F-254 plates. Bands were visualized using ultraviolet light and stained using Hanessian's stain (cerium molybdate). Flash chromatography was performed using a Teledyne-Isco CombiFlash RF 200 with UV/vis detection (254 nm, 280 nm). $^{1}$H NMR spectra were recorded on a 400 MHz Varian spectrometer using an AutoX PFG probe at 298 K; residual solvent peaks were used as internal references: DMSO (quint, δH = 2.50 ppm), $CHCl_3$ (s, δH = 7.26 ppm) or methanol (quint, δH = 3.31 ppm). $^{13}$C NMR spectra were recorded on a 400 MHz Varian spectrometer using an AutoX PFG probe at 298 K; residual solvent peaks were used as internal references: DMSO (δ = 40.50 ppm), $CHCl_3$ (δ = 77.23 ppm) or methanol (δ = 49.00 ppm). Coupling constants (J) are reported in hertz (Hz). The following abbreviations are used to describe the multiplicities: s = singlet, d = doublet, t = triplet, q = quartet, quint = quintet, sext = sextet, m = multiplet, dd = doublet-doublet, ddd = doublet-doublet-doublet, dt = doublet-triplet, dq =doublet-quartet.

#### Absorption and Fluorescence Measurements

All photophysical experiments were measured in a quartz sub-micro cuvette (10 mm path length) purchased from Starnacell Inc. Solutions were prepared in 1X PBS buffer at pH 7.4. Steady state emission scans were recorded using a PTI QuantaMaster QM-400 and absorbances were measured on a Shimadzu UV-1700 Pharmaspec spectrometer. Quantum yield measurements were performed the comparative method of Williams *et al.* and measured in duplicate, at minimum.[2] Coumarin 153 and Quinine sulfate in EtOH was used as a reference standard for quantum yield measurements. All measurements were taken with an absorbance range of 0.01-0.1. Subsequent dilutions were performed stepwise in order to obtain a minimum of five absorbance and emission spectra for quantum yield determinations. Extinction coefficient measurements were performed on samples whose concentration was determined by 1H NMR using anisole as an internal standard. Measurements were taken with an absorbance range of 0.1-1. Representative data from quantum yield measurements is plotted below. Quantum yield and extinction coefficient determinations were obtained using the following equations:

$$\Phi_{\mathrm{X}} = \Phi_{\mathrm{Std}}\left(\frac{\mathrm{Slope}_{\mathrm{X}}}{Slope_{\mathrm{Std}}}\right)\left(\frac{\eta_{\mathrm{X}}{}^{2}}{\eta_{\mathrm{Std}}{}^{2}}\right)$$

**Plasmids**

These plasmids, FM5-G3BP1-mCherry, FM5-DCP1A-mCherry and FM5-DDX6-mCherry were generous gifts from Clifford P.Brangwynne lab, Princeton University.

**Cell culture and metabolic labeling**

HeLa cells were cultured at 37 °C in a humidified atmosphere with 5% $CO_2$ in DMEM (Life Technologies) supplemented with 10% fetal bovine serum (Atlanta), 1x penicillin-streptomycin (Life Technologies) and 2 mM L-glutamine (Life Technologies). For nucleoside labeling experiments with WT HeLa cells, cells were seeded at $0.5 \times 10^6$ cells in MatTek glass bottom petri dishes. 24 hours later, fresh medium containing 500 μM PyrroloC, 500 μM tC, 500 μM $^{\mathrm{MeO}}$tC, or DMSO was added and incubated for 12 hours. For $^{\mathrm{DEA}}$tC treatment, 200 μM was incubated with HeLa cells for 6 hours because of the high toxicity of $^{DEA}$tC. For treatments with ProTide fluorescent nucleosides, 100 μM ProTide PyrroloC, 100 μM ProTide tC or 100 μM $^{DEA}$tC was added and incubated for 12 hours.

For nucleoside labeling experiments in Flp-In HeLa cells overexpressing UCK2, HeLa cells were seeded at 0.4 × $10^6$ cells per well in 35 mm glass bottom petri dishes. After 16 hours, cells were induced for 16 hours with 1 μg/mL tetracycline, and then treated with 500 μM fluorescent nucleosides for 12 hours. Control samples without tetracycline induce were also treated with fluorescent nucleosides parallelly. For cotreatments with RNA or DNA synthesis inhibitor, induced Flp-In Hela UCK2 was pretreated with 2 μM actinomycin D (ActD) or 10 mM hydroxyurea for 30 mins, and then incubated in medium containing fluorescent nucleoside and inhibitor for 6 hours.

**Analyses of fluorescent nucleoside incorporation by LC-MS/MS**

For transient transfection HeLa cells with UCK2 plasmids, HeLa cells were seeded at 0.4 × $10^6$ cells per well in six-well plate. 24 hours later, cells were transfected with pCDNA5/FRT/TO-UCK2WT plasmid (2 μg) and Lipofectamine 2000 reagent (Invitrogen, 6 μL) or pCDNA5/FRT/TO-UCK2Y65G plasmid (2 μg) and Lipofectamine 2000 reagent (Invitrogen, 6 μL) according to the manufacturer’s instructions. 12 hours after transfection, cells were treated with 500 μM fluorescent nucleosides for another 12 hours. The cells were then harvested and total RNA was extracted using Trizol reagent (Invitrogen) according to the manufacturer’s instructions.

Each RNA sample (10 μg) was digested with nuclease P1 (Wako Chemicals, 2U) in 30 μL of buffer (7 mM NaOAc, 0.4 mM $ZnCl_2$, pH 5.2) at 37 °C for 2 hours. The mixture was dephosphorylated with Antarctic Phosphatase (NEB, 2 μL) at 37 °C for 2 hours.

UHPLC-MS/MS analysis of digested RNA samples was performed on an Agilent 6470 triple quadrupole LC/MS system. The following source parameters were used for the mass spectrometry: gas temperature 175 °C, gas flow 12 L/min, nebulizer 20 psi, sheath gas temperature 325 °C, sheath gas flow 12 L/min, capillary voltage 2,500 V in the positive ion mode, capillary voltage -2,500 V in the negative ion mode. Chromatography was performed on a Thermo Scientific Hypersil GOLD aQ column (3 μm, 150 x 2.1 mm) at 36 °C using a gradient of water (containing 0.1% formic acid) and acetonitrile, at a flow rate of 0.4 mL/min: 0% ACN from 0-7 min, 10% ACN from 8-10 min, 80% ACN from 11-13 min, 0% ACN from 13.5-16.5 min. The injection volume was 1 μL. The mass transitions and retention time shown in Supplementary Table 2 were used to identify each nucleoside. Calibration curves for quantification were generated by injection of various concentrations of nucleoside standards (Figure S5).

***In vitro* enzyme assay**

Wild-type UCK2 was expressed and purified as previously reported.[1] The catalytic activity of wild-type UCK2 enzyme was tested by preparing reactions containing enzyme (500 nM), nucleoside (1 mM), KCl (100 mM), $MgCl_2$ (5 mM), NaF (15 mM), and ATP (5 mM) in Tris buffer (50 mM, pH 7.6). Reactions were incubated at 37 °C for 0.5, 1, 3, 6, 12, 18 hours and quenched by heating at 95 °C for 5 minutes. After centrifugation (17,000 *g* for 10 minutes), reaction with 5-ICyt and 5-BrCyt were analyzed by HPLC on an Agilent Poroshell 120 EC-

C18 (4 μm, 4.6×150 mm) column using a 0.1 M TEAA/MeOH gradient (0% to 1.5% methanol from 0-10 minutes, 1.5% to 2.0% methanol from 10-20 minutes, followed by 2.0% to 4.5% methanol from 20-30 minutes) at a flow rate of 1.2 mL/min. Chromatography for reactions with PyrroloC, tC, $^{MeO}$C or $^{DEA}$tC was performed using the 0.1 M TEAA/ACN gradient (0% to 15% ACN from 0-5 minutes, 15% to 40% methanol from 5-30 minutes) at a flow rate of 1.2 mL/min. Product identities were confirmed by HRMS analysis on an Agilent 6220 ESI-TOF (Supplementary Table 1).

**Docking**

Crystal structure of the UCK2-Cytidine complex was obtained from the Protein Data Bank (PDB accession 1UEJ). Explicit hydrogen atoms were added and all water molecules were then deleted. The structure of PyrroloC with UCK2 was generated using AutoDock Tools (http://autodock.scripps.edu/). In brief, Cytidine was removed, PyrroloC was prepared for docking by setting all rotatable bonds as active torsions. Next, PyrroloC was docked into UCK2 using AutoDockVina (version 1.11; Scripps Research Institute). The grid box was centered on the ligand in the active site of original crystal structure. The macromolecule molecular surface and secondary structure elements were displayed by PyMol. The interaction between UCK2 and PyrroloC was shown by the polar contacts function.

**Live cell confocal fluorescence microscopy**

Cells were seeded and cultured through the protocol aforementioned. 200 μM tC was used for labeling experiments. Cells were washed with DPBS for twice, and fresh DMEM for 5 mins. Leibovitz's L-15 Medium was used for live cell imaging. For epifluorescence microscopy imaging, DAPI channel was used for detection the fluorescent nucleosides signal. For confocal fluorescence microscopy imaging of tC labeled cells, the microscope environment was maintained at 37°C and 5% of $CO_2$. We use 405 nm laser as excitation, green fluorescent protein (GFP) filter sets for emission and 60-fold magnification objective. For each imaging, z-stacks of 11 pictures were acquired in 10 μm. All z-stacks were maximum-projected before analysis. Quantification of signal was used ImageJ. For colocalization experiments in figure 4I, single z-stack imaging result was used.

**RNA Synthesis under different serum concentration**

Flp-In HeLa cells overexpressing UCK2 were seeded at $0.4 \times 10^6$ cells per well in 35 mm glass bottom petri dishes. After 16 hours, cells were induced with 1 μg/mL tetracycline in 5% FBS DMEM medium for 16 hours. Then treated with 200 μM tC in 5% or 10% FBS DMEM medium for 1 hours. For negative control, induced cells were treated with 2 μM ActD for 1 hour first, and then cotreated with 200 μM tC for another 1 hour.

**RNA Synthesis under stress**

Induced Flp-In HeLa cells overexpressing UCK2 pretreated with 200 μM $NaAsO_2$ for 0.5 hour, and then treated with 200 μM tC together with 200 μM $NaAsO_2$ for indicated time.

**Pulse-chase experiments**

Induced Flp-In HeLa cells overexpressing UCK2 were treated with 200 μM tC for 4 hours of pulse labeling. Next, cells were washed three times in DPBS, and incubated in fresh DMEM media or 200 μM $NaAsO_2$ in DMEM media for 0.5 hour.

**Colocalization of RNA foci under stress with biomarker proteins**

Flp-In HeLa cells overexpressing UCK2 were seeded at $0.4 \times 10^6$ cells per well in 35 mm glass bottom petri dishes. After 16 hours, cells were transfected with desired constructs plasmid (100 ng) and Lipofectamine 2000 reagent (Invitrogen, 2 µL) according to the manufacturer's instructions. 5 hours after transfection, cells were induced with 1 µg/mL tetracycline for 16 hours. cells were treated with tC through the protocol aforementioned in RNA synthesis under stress.

**Cell viability assay**

Cells were plated at 2000 cells per well in 96-well plates. After 16 hours, 1 µg/mL tetracycline was used to induce expression of UCK2 gene. Following 12 hours of induction, cells were treated with PyrroloC (10-1000 µM) or tC (10-1000 µM) for 24 hours. CellTiter 96 Aqueous One Solution Cell Proliferation Assay (MTS) (Promega) was performed according to the manufacturer's instructions. Absorbance was measured at 490 nm, and cell viability was calculated as the ratio of absorbance of PyrroloC or tC treated cells to that of untreated cells.

**Supplementary Table 1**: quantum yield and extinction coefficients of the ribonucleosides.

| **Compound** | **Excitation Wavelength (nm)** | **Quantum Yield** | **Extinction Coefficient ($M^{-1}$ $cm^{-1}$)** |
|---|---|---|---|
| tC | 377 | 0.17 | 5351 |
| pyrrolo-C | 337 | 0.54 | 2905 |
| $^{MeO}$tC | 379 | 0.015 | 5015 |
| $^{DEA}$tC | 395 | 0.006 | 2416 |

**Supplementary Table 2.** Dynamic multiple reaction monitoring (DMRM) parameters of for quantification nucleosides by LC-QQQ-MS.

| Nucleoside | Fragmentor Energy (V) | Collision Energy (V) | Parent Ion $[M+H]^+$ | Product Ion $[M+H]^+$ | Retention time (min) | Source/Vendor |
|---|---|---|---|---|---|---|
| A | 100 | 20 | 268.1 | 136.1 | 3.7 | Sigma |
| G | 80 | 13 | 284.1 | 152.1 | 4.3 | Sigma |
| C | 70 | 14 | 244.1 | 112.1 | 1.7 | Sigma |
| tC | 70 | 14 | 350.3 | 218.2 | 13.6 | Self-synthesized |
| PyrroloC | 70 | 14 | 282.2 | 150.1 | 11.9 | Self-synthesized |
| $^{DEA}$tC | 70 | 14 | 421.5 | 289.3 | 13.2 | Self-synthesized |
| $^{MeO}$tC | 70 | 14 | 380.4 | 248.3 | 13.8 | Self-synthesized |
| U | 70 | 7 | 245.1 | 113.1 | 2.3 | Sigma |

**Supplementary Table 3.** Nucleoside concentration (ng/mL) in total RNA of HeLa cells transfected with WT or mutant versions of UCK2. Experiments were performed in triplicate. Nucleosides marked with † were quantified in 200 ng of RNA. All others were quantified in 1 ng of RNA.

| Transfection | Treatment | C | U | A | G | tC† | PyC† |
|---|---|---|---|---|---|---|---|
| – | tC | 192.1 | 87.7 | 140.2 | 360.1 | 5.36 | 0 |
| – | tC | 210.9 | 95.8 | 154.67 | 397.1 | 4.91 | 0 |
| – | tC | 135.0 | 64.7 | 60.47 | 57.4 | 5.74 | 0 |
| UCK2 WT | tC | 125.98 | 69.6 | 178.7 | 368.0 | 30.1 | 0 |
| UCK2 WT | tC | 141.4 | 76.4 | 151.4 | 302.4 | 30.7 | 0 |
| UCK2 WT | tC | 97.6 | 55.9 | 207.7 | 361.5 | 28.5 | 0 |
| UCK2 Y65G | tC | 121.6 | 60.0 | 172.2 | 251.7 | 25.9 | 0 |
| UCK2 Y65G | tC | 117.8 | 64.6 | 187.2 | 213.4 | 27.5 | 0 |
| UCK2 Y65G | tC | 120.5 | 65.5 | 214.3 | 228.0 | 26.1 | 0 |
| – | PyrroloC | 118.9 | 56.8 | 150.3 | 323.3 | 0 | 1.07 |
| – | PyrroloC | 123.0 | 62.9 | 142.8 | 414.1 | 0 | 1.16 |
| – | PyrroloC | 128.0 | 67.0 | 175.2 | 401.4 | 0 | 1.04 |
| UCK2 WT | PyrroloC | 98.9 | 58.2 | 200.5 | 384.7 | 0 | 101.4 |
| UCK2 WT | PyrroloC | 120.8 | 65.4 | 189.2 | 382.5 | 0 | 114.1 |
| UCK2 WT | PyrroloC | 94.6 | 55.3 | 170.2 | 351.4 | 0 | 98.9 |
| UCK2 Y65G | PyrroloC | 392.4 | 103.7 | 400.2 | 521.5 | 0 | 137.7 |
| UCK2 Y65G | PyrroloC | 47.1 | 31.1 | 121.4 | 245.6 | 0 | 38.1 |
| UCK2 Y65G | PyrroloC | 65.7 | 41.1 | 115.3 | 225.7 | 0 | 49.5 |

**Supplementary Table 4.** Nucleoside concentration (ng/mL) in total RNA of FlpIn HeLa cells expressing WT or mutant versions of UCK2. Experiments were performed in triplicate. Nucleosides marked with † were quantified in 200 ng of RNA. All others were quantified in 1 ng of RNA.

| Induction | Treatment | C | U | A | G | tC† | Py C† |
|---|---|---|---|---|---|---|---|
| – | tC | 128.9 | 70.1 | 140.8 | 266.1 | 15.7 | 0 |
| – | tC | 118.4 | 61.1 | 108.7 | 243.1 | 15.5 | 0 |
| – | tC | 127.5 | 66.9 | 120.8 | 257.4 | 15.2 | 0 |
| UCK2 WT | tC | 88.3 | 57.4 | 128.7 | 202.0 | 44.2 | 0 |
| UCK2 WT | tC | 100.4 | 59.2 | 115.4 | 202.5 | 42.1 | 0 |
| UCK2 WT | tC | 100.5 | 58.7 | 168.7 | 266.5 | 43.1 | 0 |
| UCK2 Y65G | tC | 133.6 | 81.7 | 118.0 | 241.7 | 103.9 | 0 |
| UCK2 Y65G | tC | 140.5 | 80.8 | 111.8 | 230.4 | 105.5 | 0 |
| UCK2 Y65G | tC | 128.5 | 78.2 | 214.3 | 228.0 | 99.8 | 0 |
| – | Pyrrolo C | 128.9 | 71.1 | 120.7 | 328.3 | 0 | 4.65 |
| – | Pyrrolo C | 111.9 | 60.0 | 131.8 | 400.2 | 0 | 4.22 |
| – | Pyrrolo C | 112.4 | 62.4 | 128.6 | 383.2 | 0 | 4.59 |
| UCK2 WT | Pyrrolo C | 116.1 | 63.6 | 100.5 | 383.7 | 0 | 143.5 |
| UCK2 WT | Pyrrolo C | 105.1 | 65.5 | 109.2 | 326.5 | 0 | 141.9 |
| UCK2 WT | Pyrrolo C | 112.3 | 64.6 | 120.2 | 313.6 | 0 | 144.4 |
| UCK2 Y65G | Pyrrolo C | 142.1 | 86.1 | 125.9 | 255.2 | 0 | 164.3 |
| UCK2 Y65G | Pyrrolo C | 136.4 | 77.4 | 108.5 | 222.4 | 0 | 166.8 |
| UCK2 Y65G | Pyrrolo C | 150.2 | 89.5 | 134.5 | 260.2 | 0 | 177.5 |

**Supplementary Table 5.** $^{DEA}$tC and $^{MeO}$tC nucleoside concentration (ng/mL) in total RNA of FlpIn HeLa cells expressing WT or mutant versions of UCK2. Experiments were performed in triplicate. Nucleosides marked with † were quantified in 200 ng of RNA. All others were quantified in 1 ng of RNA.

| Induction | Treatment | C | U | A | G | $^{DEA}$tC† | $^{MeO}$tC † |
|---|---|---|---|---|---|---|---|
| – | – | 246.6 | 114.6 | 167.1 | 633.7 | 0.0 | 0.0 |
| – | – | 252.7 | 114.7 | 168.8 | 635.5 | 0.0 | 0.0 |
| – | – | 257.1 | 117.8 | 167.9 | 636.5 | 0.0 | 0.0 |
| – | $^{DEA}$tC | 246.2 | 114.8 | 161.7 | 615.7 | 0.0 | 0.0 |
| – | $^{DEA}$tC | 244.0 | 115.5 | 169.9 | 630.0 | 0.0 | 0.0 |
| – | $^{DEA}$tC | 271.4 | 127.5 | 218.6 | 785.3 | 0.0 | 0.0 |
| UCK2 WT | $^{DEA}$tC | 241.3 | 108.8 | 162.6 | 666.3 | 0.0 | 0.0 |
| UCK2 WT | $^{DEA}$tC | 268.7 | 126.3 | 174.5 | 663.7 | 0.0 | 0.0 |
| UCK2 WT | $^{DEA}$tC | 227.3 | 103.7 | 147.7 | 561.9 | 0.0 | 0.0 |
| – | $^{MeO}$tC | 247.5 | 115.2 | 160.0 | 610.1 | 0.0 | 3.64 |
| – | $^{MeO}$tC | 249.1 | 115.8 | 197.5 | 705.3 | 0.0 | 4.16 |
| – | $^{MeO}$tC | 236.4 | 109.8 | 148.4 | 580.5 | 0.0 | 3.84 |
| UCK2 WT | $^{MeO}$tC | 227.6 | 107.2 | 166.3 | 645.0 | 0.0 | 12.2 |
| UCK2 WT | $^{MeO}$tC | 236.1 | 109.7 | 142.5 | 571.1 | 0.0 | 11.8 |
| UCK2 WT | $^{MeO}$tC | 228.2 | 104.7 | 165.3 | 647.7 | 0.0 | 12.3 |

**Supplementary Table 6.** ESI-MS of product generated by UCK2 phosphorylation of PyrroloC, tC, $^{MeO}$tC, 5-BrCyt and 5-ICyt.

| | Calculated $[M\text{-}H]^-$(m/z) | Detected $[M\text{-}H]^-$ (m/z) |
|---|---|---|
| PyrroloC -$PO_4$ ($C_{12}H_{16}N_3O_8P$) | 360.0602 | 360.0604 |
| tC-$PO_4$-$PO_4$ ($C_{15}H_{16}N_3O_8PS$) | 428.0323 | 428.0326 |
| $^{MeO}$tC-$PO_4$ ($C_{16}H_{18}N_3O_9PS$) | 458.0423 | 458.0434 |
| 5-BrCyt-$PO_4$ ($C_9H_{13}N_3O_8PSBr$) | 399.5991 | 399.5993 |
| 5-ICyt-$PO_4$ ($C_9H_{13}N_3O_8PSI$) | 477.9412 | 477.9415 |

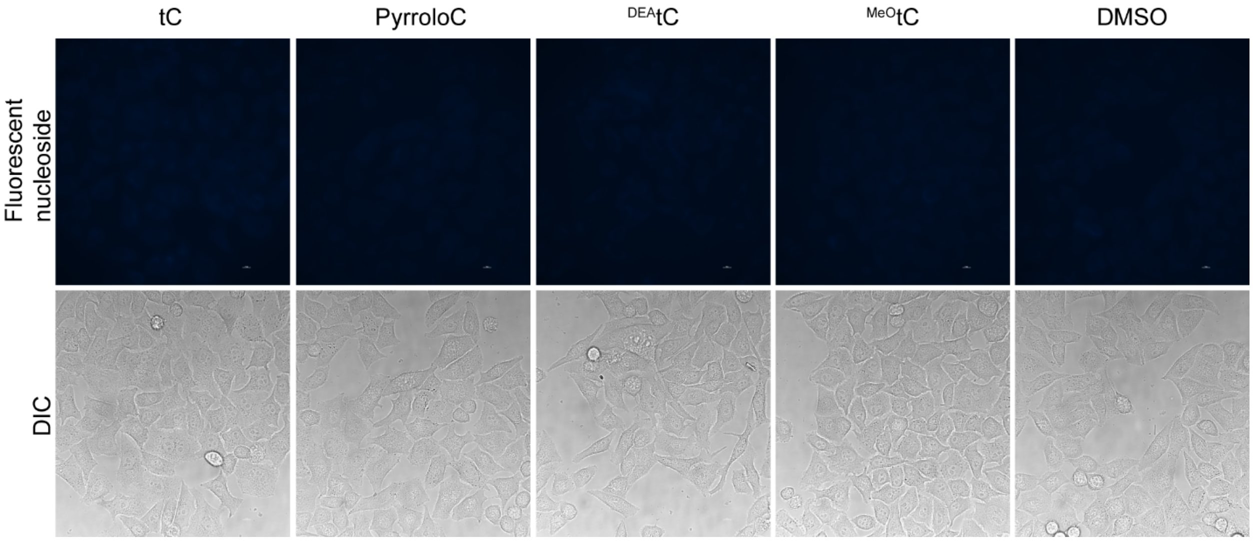


**Supplementary Figure 1.** Labeling of wt HeLa cells with 500 μM fluorescent nucleosides. For $^{DEA}$tC, 200 μM was used for live cell treatment.

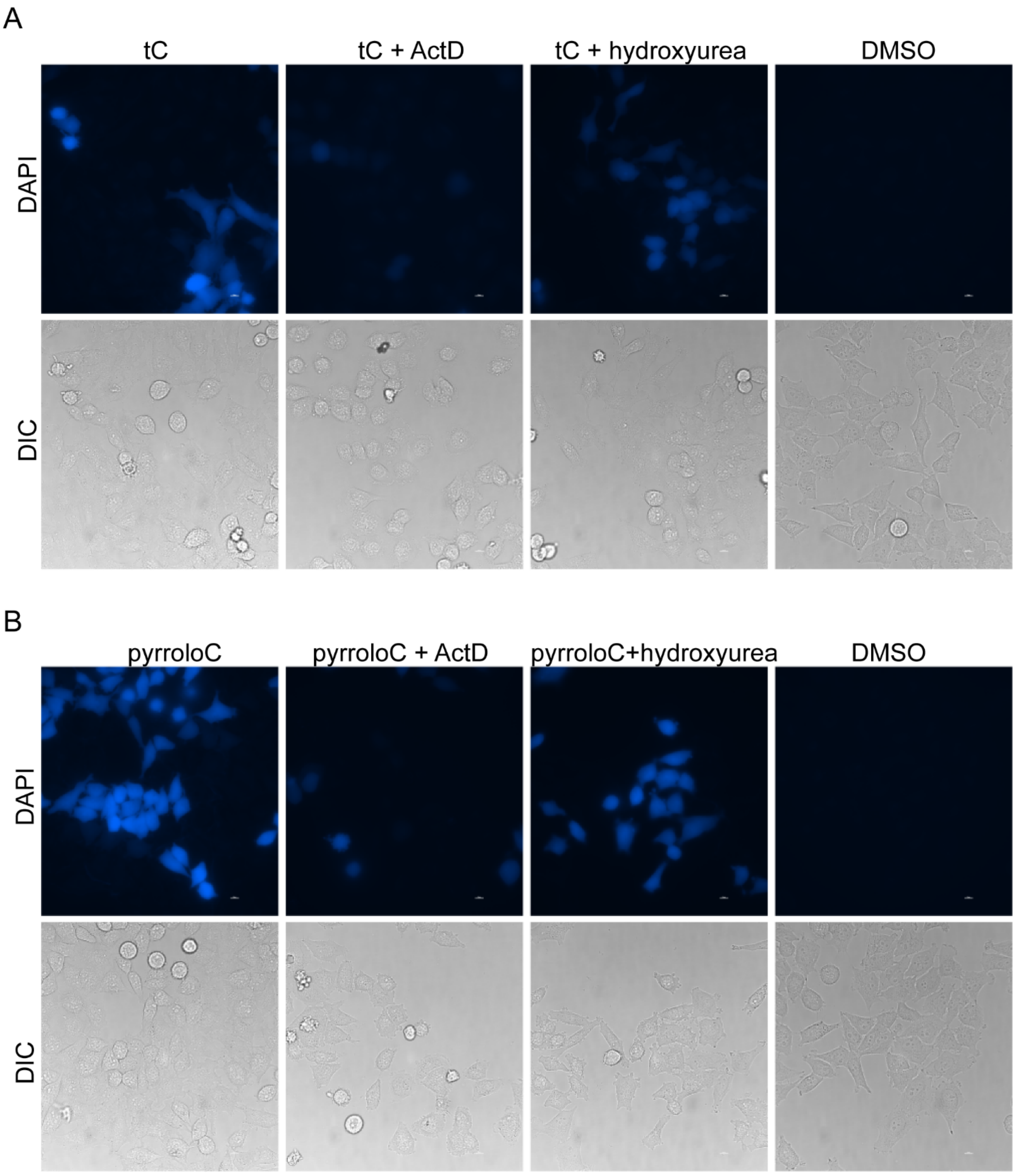


**Supplementary Figure 2.** Labeling of HeLa cells expressing WT UCK2 with tC or PyrroloC in the presence of the RNA polymerase inhibitor actinomycin D or DNA synthesis inhibitor hydroxyurea. (A) Cells were treated with 200 μM tC in the presence or absence of the indicated concentrations of Actinomycin D for 5 hr. (B) Cells were treated with 200 μM PyrroloC in the presence or absence of the indicated concentrations of Actinomycin D for 5 hrs.

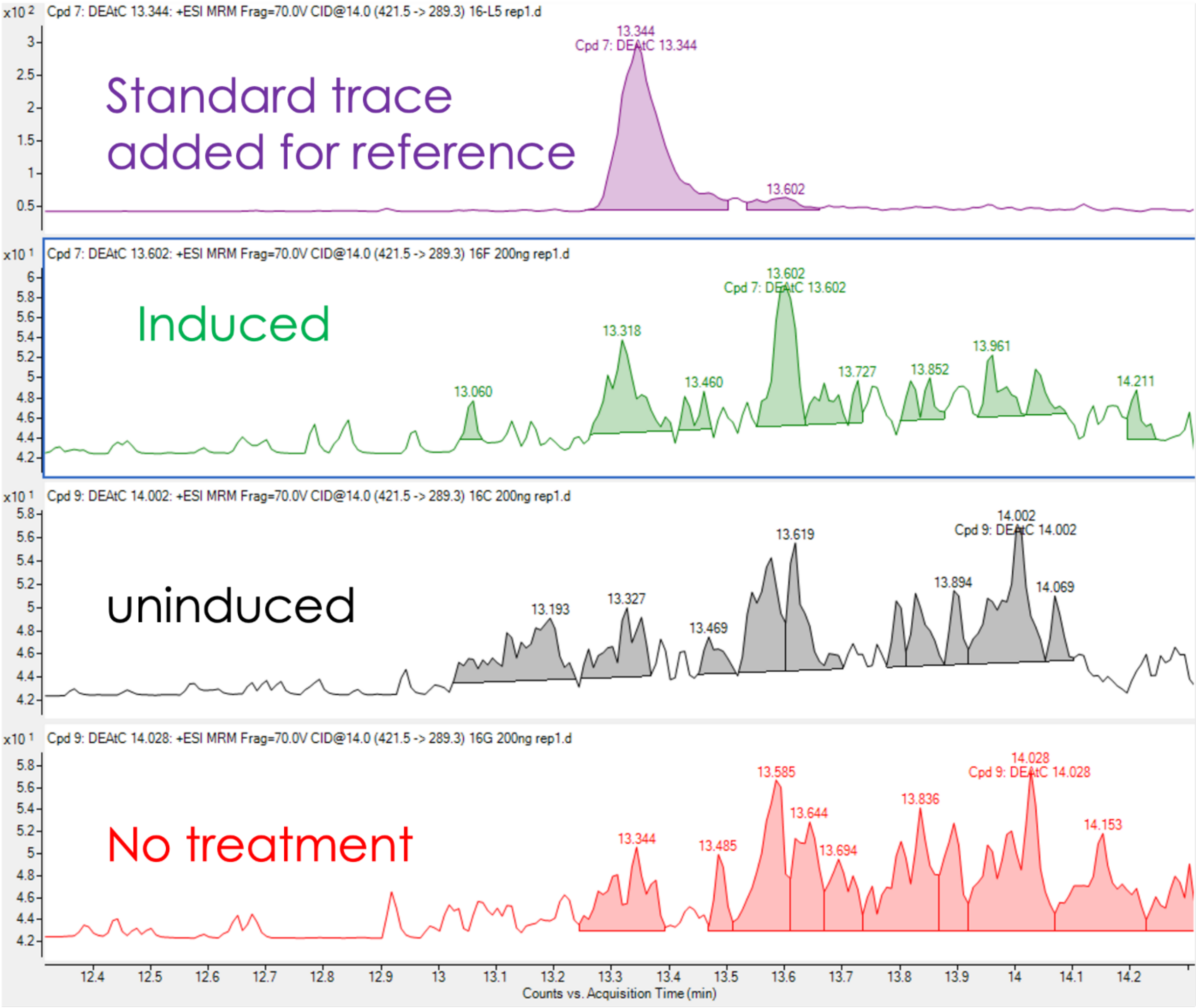


**Supplementary Figure 3.** Induced/ uninduced HeLa Flpin UCK2 cells were treated with $^{DEA}$tC. Incorporation of $^{DEA}$tC was detected by LCMS/MS, chromatographic trace of $^{DEA}$tC was shown.

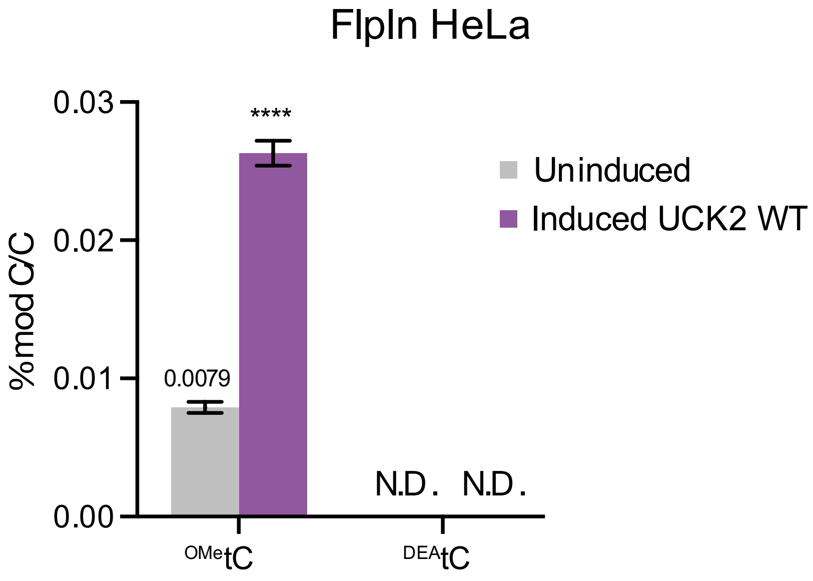


**Supplementary Figure 4.** Induced/ uninduced HeLa Flpin UCK2WT cells were treated with $^{MeO}$tC or $^{DEA}$tC. Incorporation of $^{MeO}$tC or $^{DEA}$tC was detected by LCMS/MS. N.D. means not detected.

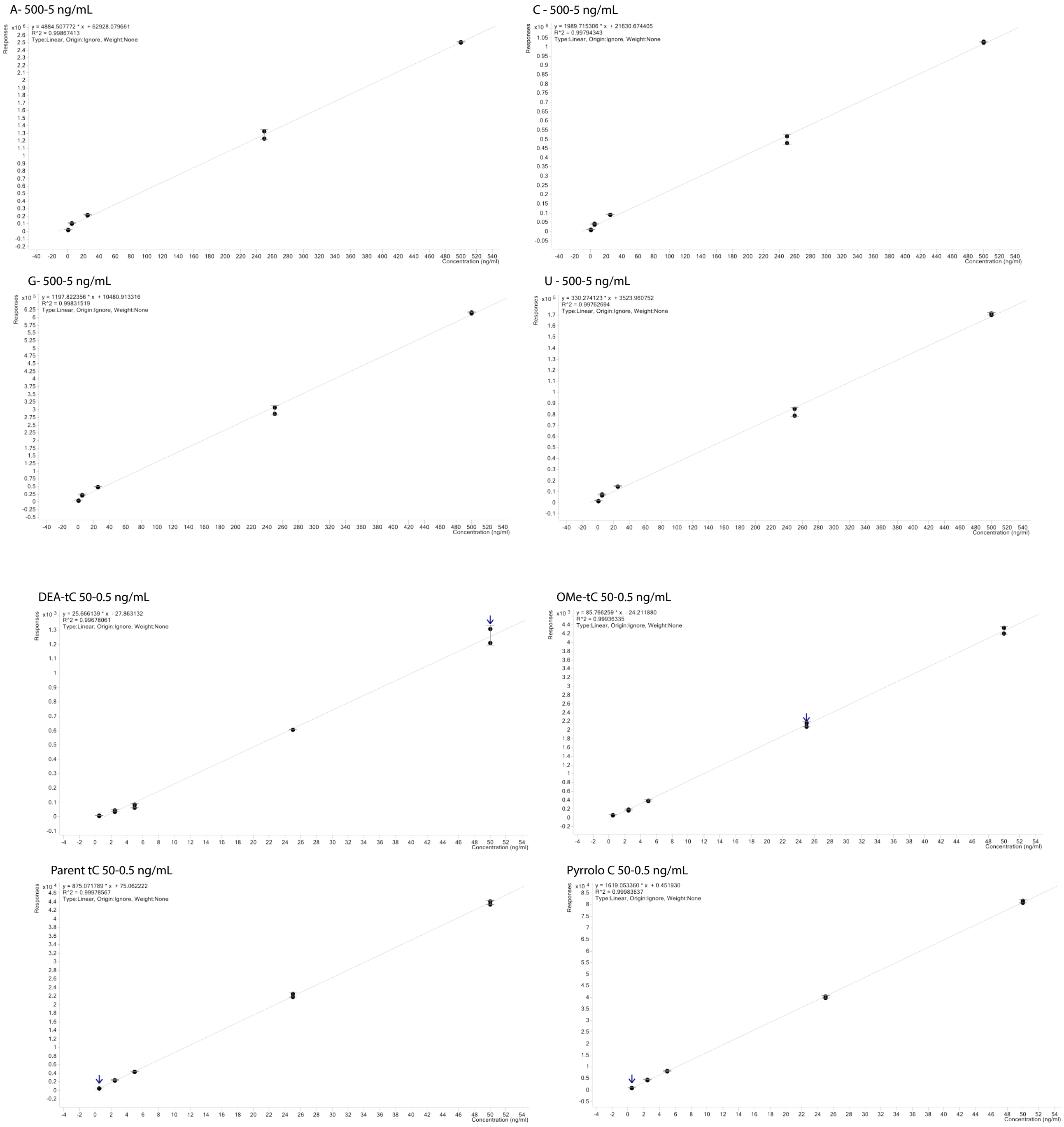


**Supplementary Figure 5**. Representative standard curves for native and modified nucleosides analyzed by quantitative LC-MS/MS

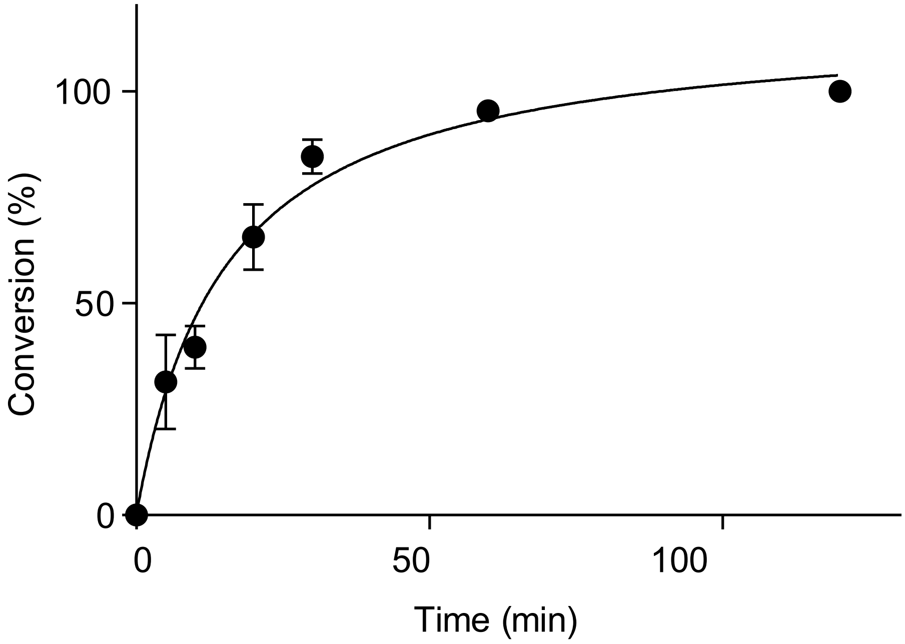


**Supplementary Figure 6** Quantification of PyrroloC phosphorylation conversion by WT UCK2. Reactions containing UCK2, nucleoside, and ATP in buffer were run for indicated time at 37 °C and subsequently analyzed by RP-HPLC. Data represent the mean +/- s.d. (n=3).

**(A)**

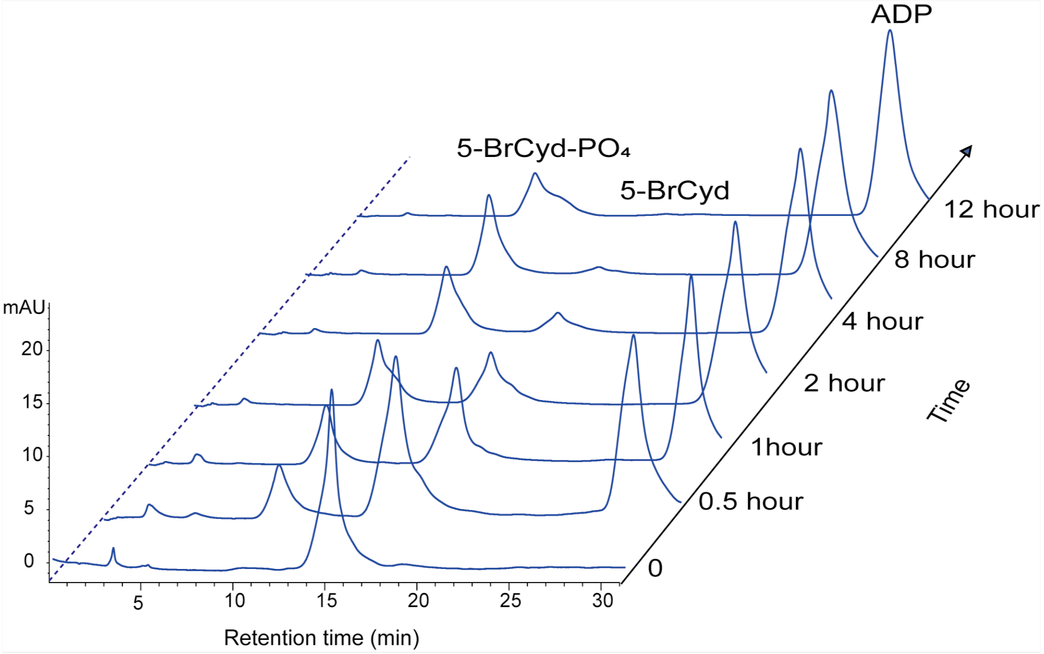


**(B)**

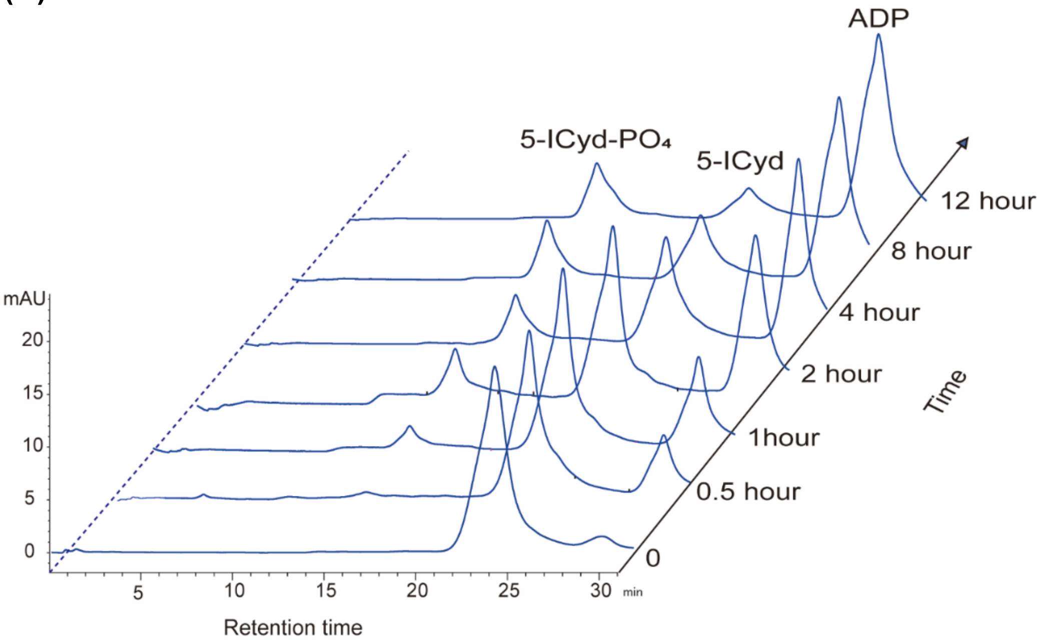


**Supplementary Figure 7.** RP-HPLC chromography of 5-BrCyd(A), 5-ICyd(B) phosphorylation by UCK2 enzymes. Reactions containing WT UCK2, cytidine analogue, and ATP in buffer were run at 37 °C for indicated time and subsequently analyzed by RP-HPLC. The retention times of nucleotide standards are indicated.

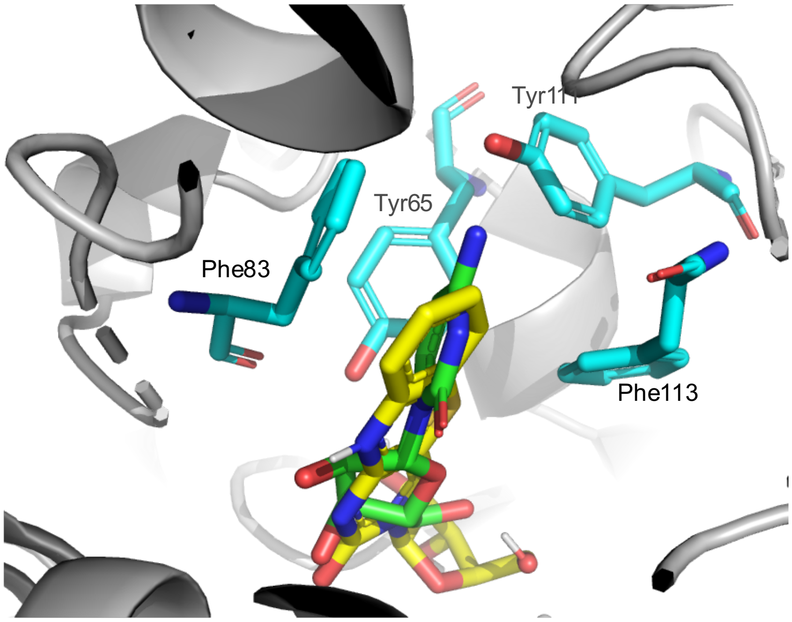


**Supplementary Figure 8.** Structural model of tC (golden) bound to UCK2 (PDB: 1UEJ).

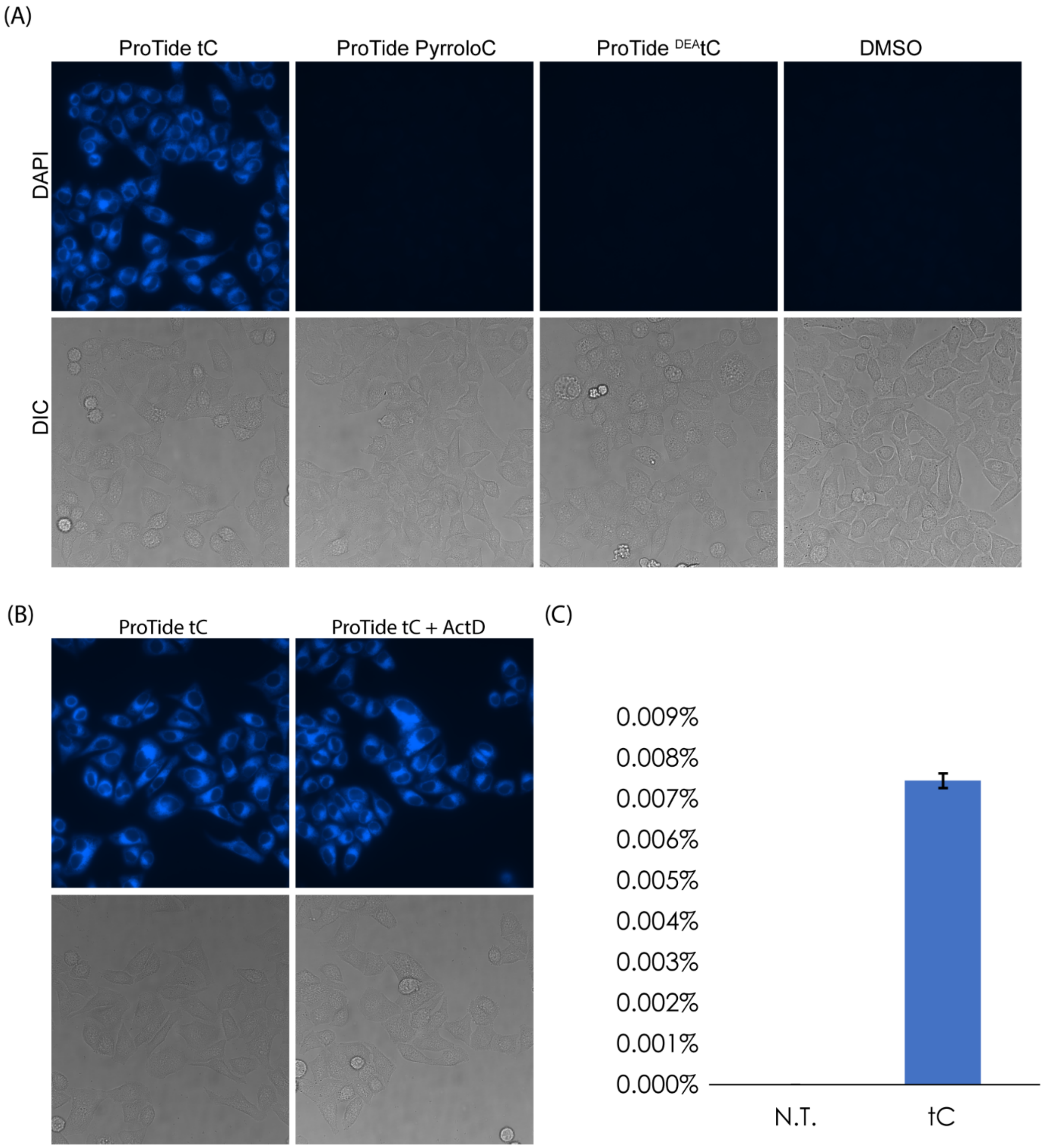


**Supplementary Figure 9.** Labeling of wt HeLa cells with 100 μM ProTide fluorescent nucleosides. (A) wt HeLa cells were treated with ProTide version of tC, PyrroloC, or $^{DEA}$tC. (B) wt HeLa cells were cotreated with ProTide and 2 μM Actinomycin D for 5 hours. (C) wt HeLa cells were treated with 100 μM ProTide tC for overnight, total RNA was extracted and digested, and tC incorporation was quantified by LCMS/MS.

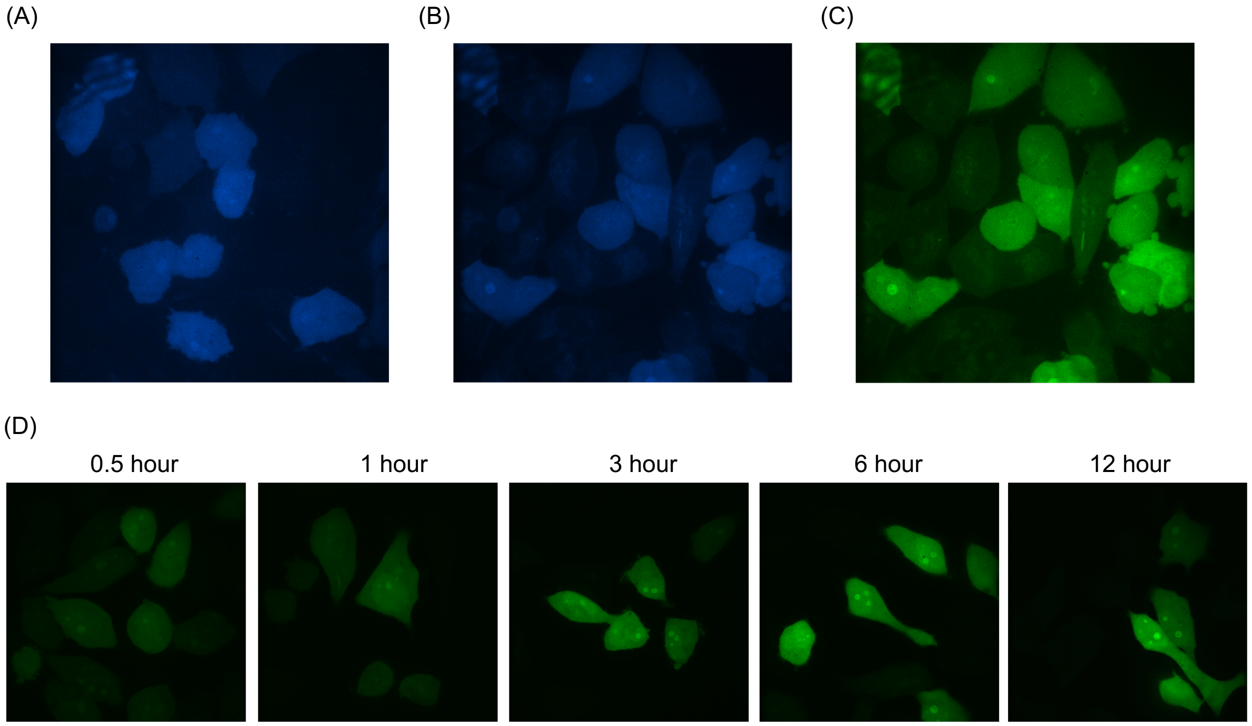


**Supplementary Figure 10**. Optimization of confocal microscope imaging. (A) 200 μM pyrroloC or (B) 200 μM tC treated HeLa UCK2 cells for 6 hours, image result was collected using 405 nm laser as excitation, 460±25 nm filter and 1 second exposure time. (C) View field of (B) was also imaged using 405 nm laser as excitation, 535+20 nm filter and 0.4 second exposure time. (D) Time-course analysis of cellular RNA lalbeling with 200 μM tC.

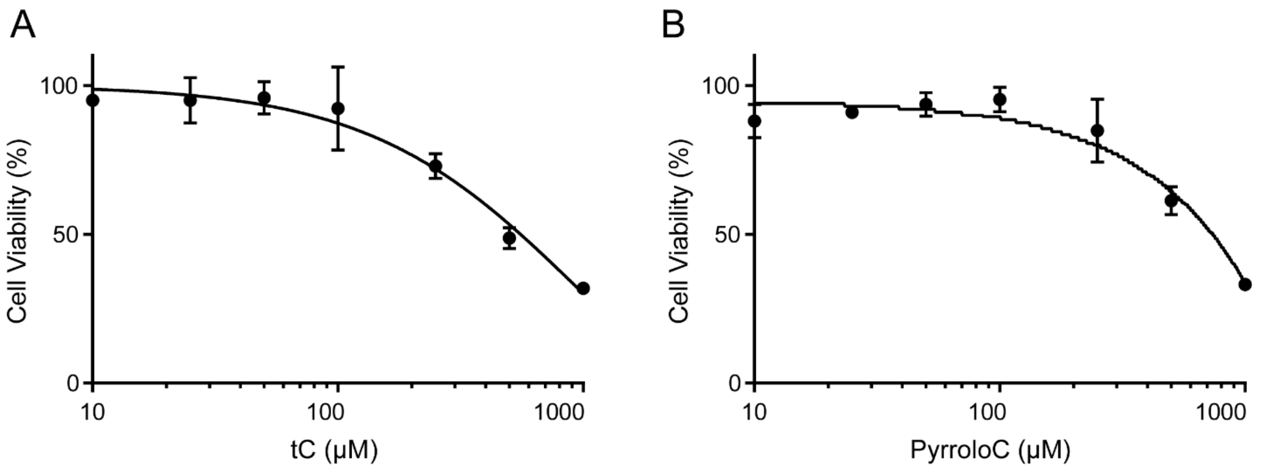


**Supplementary Figure 11**. HeLa cells expressing WT UCK2 viability was measured using an MTS-based assay under tC or PyrroloC treatment. Data represent the mean +/- s.d. (n=3).

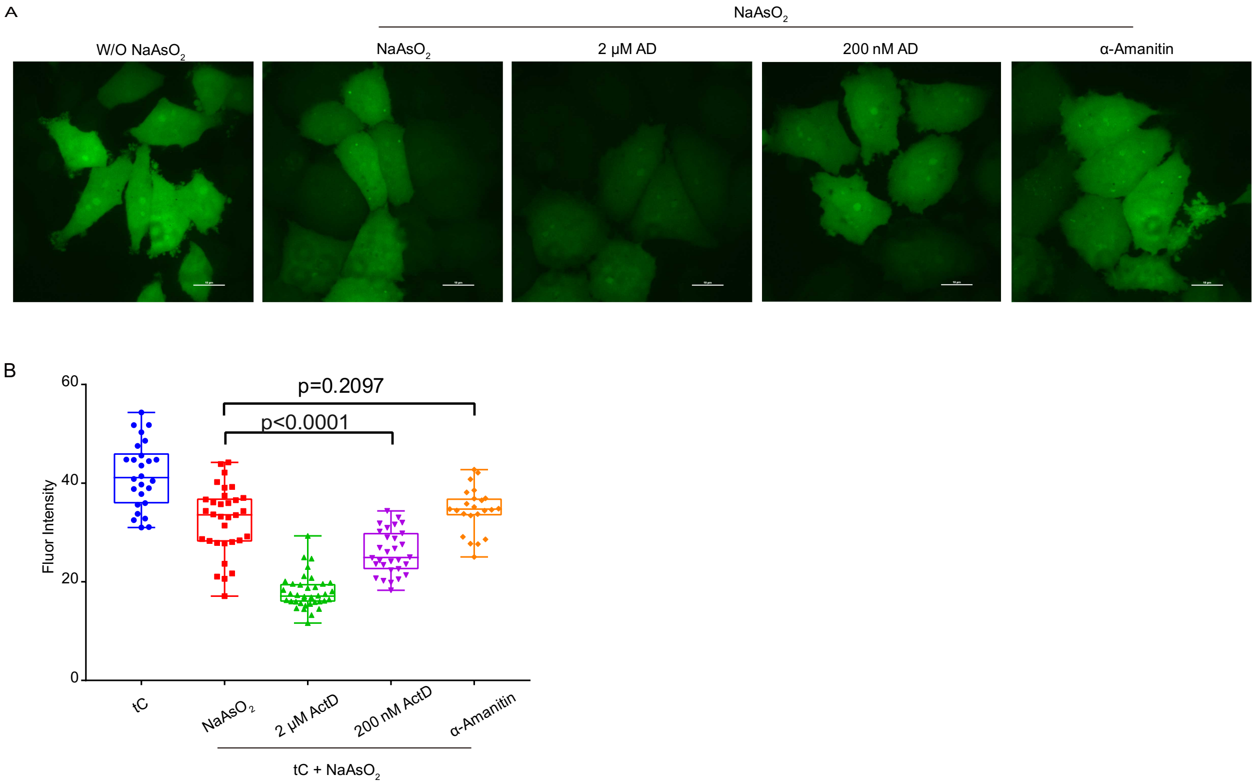


**Supplementary Figure 12.** Labeling of UCK2-expressing HeLa cells with tC in the presence of ActD or α-amanitin under stress condition. (A) Cells were treated with 200 μM tC for 1 hour together with 200 μM $NaAsO_2$ and indicated RNA polymerase inhibitor in medium. (B) Fluorescence quantification of cells treated in (A). Error bars indicate mean ± s.d. unpaired two-sided t-test, n= 25 measurements from three independent replicates.

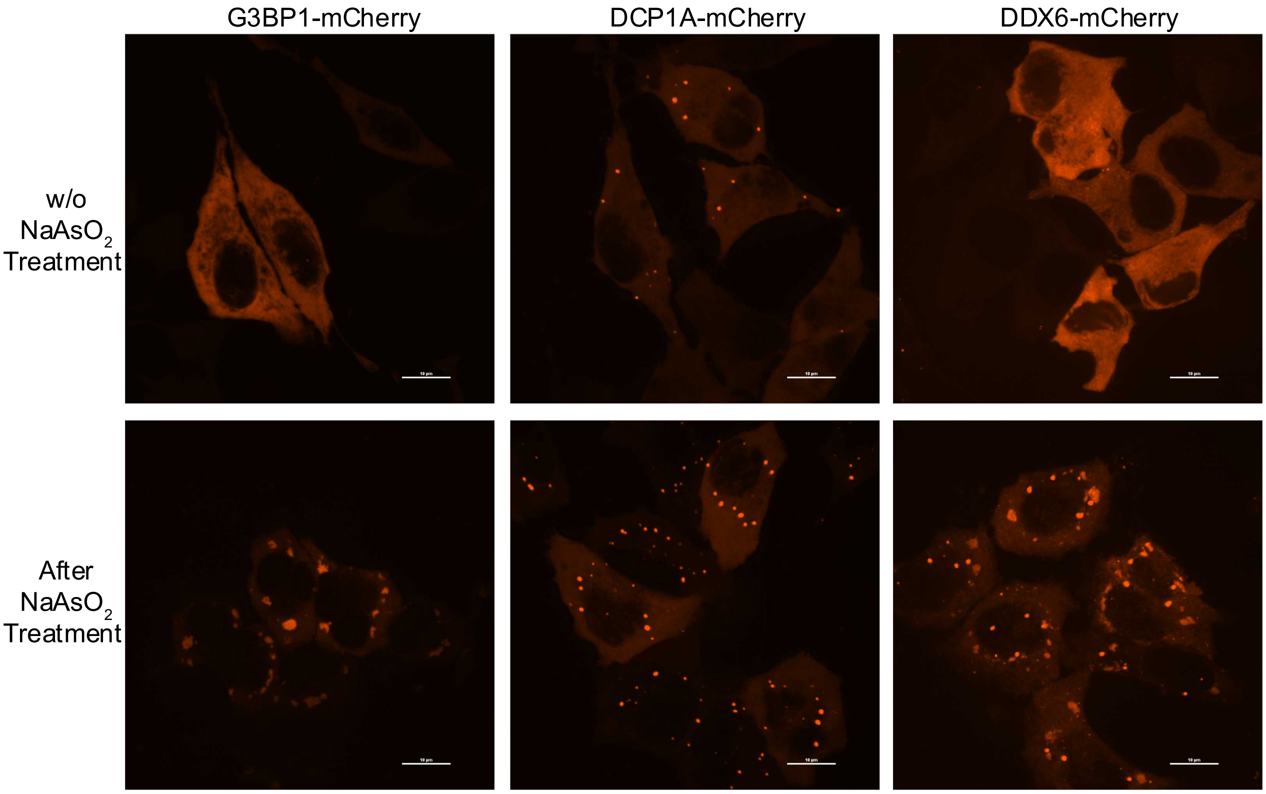


**Supplementary Figure 13.** The localization of G3BP1, DCP1A and DDX6 under stress condition. UCK2-expressing HeLa cells were transiently transfected with G3BP1-mCherry, DCP1A-mCherry or DDX6-mCherry respectively. After overnight, cells were incubated with 200 μM $NaAsO_2$ for 1.5 hour for tress condition.

### Chemical Synthesis

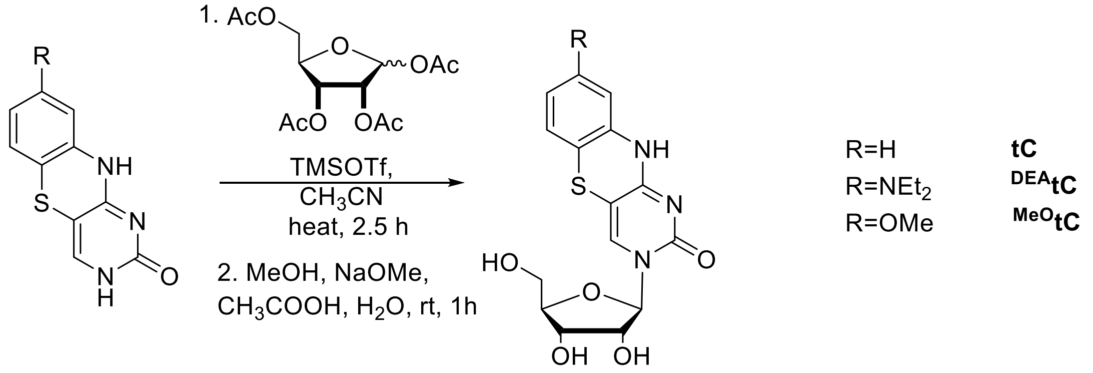


**Scheme S1. Synthesis of tC, $^{DEA}$tC and $^{MeO}$tC ribonucleosides**

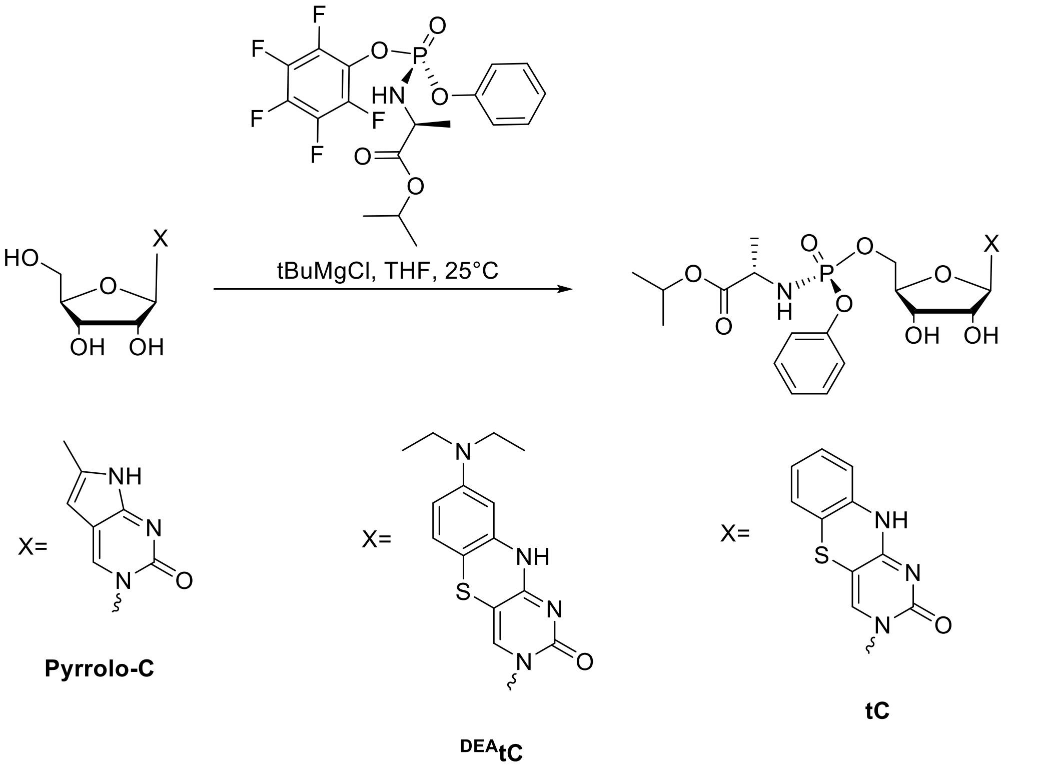


**Scheme S2: Synthesis of ProTides**

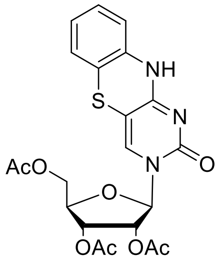


**2,3,5-tri-O-acetyl-tC-ribonucleoside.** The tC nucleobase (100 mg, 0.46 mmol) was suspended in anhydrous $CH_3CN$ (2 mL), and N,O-bis(trimethylsilyl)acetamide (168 µL, 0.69 mmol) was added. The reaction mixture was heated at 85°C under $N_2$ for 20 min and then allowed to cool to room temperature. 1,2,3,5-Tetra-O-acetyl-*β*-D-ribofuranose (175 mg, 0.55 mmol) and trimethylsilyl trifluoromethanesulfonate (100 µL, 0.55 mmol) were added. The reaction mixture was heated at 85°C for 2.5 h and allowed to cool to room temperature. The reaction progress was monitored with TLC (10% methanol in DCM). The reaction mixture was then extracted with 5% $NaHCO_3$ solution (25 mL) and $CH_2Cl_2$ (2 × 25 mL). After drying over anhydrous $Na_2SO_4$, the solvent was removed by rotary evaporation and the product was purified by flash chromatography on a Teledyne-ISCO CombiFlash Rf 200 (5% hexanes in EtOAc), yielding the semi-pure product as a yellow-orange oil that was carried through into deacetylation without further purification.

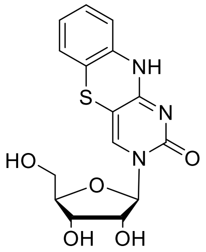


**tC-Ribonucleoside.** The semi-pure 2,3,5-Tri-O-acetyl-tC-ribonucleoside was dissolved in dry MeOH (3 mL) under $N_2$, and NaOMe (25 wt % in MeOH, 200 µL) was added. The reaction progress was monitored with TLC (10% methanol in DCM). After stirring for 1 h at room temperature, acetic acid (150 µL) was added, and the solvent removed by rotary evaporation.

The crude product was dissolved in dichloromethane and purified by flash chromatography on a Teledyne-ISCO CombiFlash Rf 200 (20% MeOH in DCM), yielding the pure product as a yellow solid (105 mg, 66% over two steps). $^{1}H$ NMR (500 MHz, DMSO-$d_6$) δ 7.95 (s, 1H), 7.07 (d, $J$ = 7.5 Hz, 2H), 6.96 – 6.90 (m, 2H), 5.74 (s, 1H), 5.72 (d, $J$ = 3.2 Hz, 1H), 3.96 (d, $J$ = 4.6 Hz, 2H), 3.86 (d, $J$ = 18.1 Hz, 1H), 3.70 (d, $J$ = 12.6 Hz, 1H), 3.57 (m, 1H); HRMS (ESI) m/z calcd for $C_{15}H_{16}N_3O_5S^+$ $[M+H]^+$ 350.0805, found 380.0806

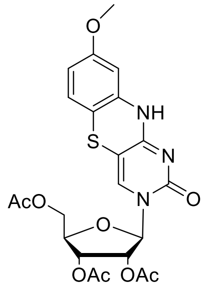


**2,3,5-tri-O-acetyl-$^{MeO}$tC-ribonucleoside.** The $^{MeO}$tC nucleobase (108 mg, 0.44 mmol) was suspended in anhydrous $CH_3CN$ (2 mL), and N,O-bis(trimethylsilyl)acetamide (160 µL, 0.65 mmol) was added. The reaction mixture was heated at 85°C under $N_2$ for 20 min and then allowed to cool to room temperature. 1,2,3,5-Tetra-O-acetyl-*β*-D-ribofuranose (166 mg, 0.52 mmol) and trimethylsilyl trifluoromethanesulfonate (95 µL, 0.52 mmol) were added. The reaction mixture was heated at 85°C for 2.5 h and allowed to cool to room temperature. The reaction progress was monitored with TLC (10% methanol in DCM). The reaction mixture was then extracted with 5% $NaHCO_3$ solution (25 mL) and $CH_2Cl_2$ (2 × 25 mL). After drying over anhydrous $Na_2SO_4$, the solvent was removed by rotary evaporation and the product was purified by flash chromatography on a Teledyne-ISCO CombiFlash Rf 200 (5% hexanes in EtOAc), yielding the semi-pure product as a yellow-orange oil that was carried through into deacetylation without further purification.

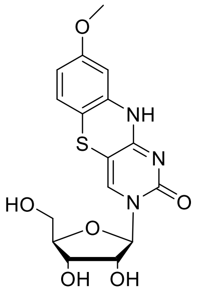

**$^{MeO}$tC-Ribonucleoside.** The semi-pure 2,3,5-Tri-O-acetyl-$^{MeO}$tC-ribonucleoside (92 mg, 0.18 mmol) was dissolved in dry MeOH (2 mL) under $N_2$, and NaOMe (25 wt % in MeOH, 200 µL) was added. The reaction progress was monitored with TLC (10% methanol in DCM). After stirring for 1 h at room temperature, acetic acid (150 µL) was added, and the solvent removed by rotary evaporation. The crude product was dissolved in dichloromethane and purified by flash chromatography on a Teledyne-ISCO CombiFlash Rf 200 (20% MeOH in DCM), yielding the pure product as a yellow solid (40 mg, 24%). 1H NMR (400 MHz, Methanol-$d_4$) δ 7.84 (s, 1H), 7.43 (s, 1H), 6.80 (d, J = 8.5 Hz, 1H), 6.48 (dd, J = 8.5, 2.5 Hz, 1H), 6.45 (d, J = 2.5 Hz, 1H), 5.75 (d, J = 3.5 Hz, 1H), 4.13 (t, J = 5.5 Hz, 1H), 4.09 (dd, J = 5.3, 3.5 Hz, 1H), 4.04 (dt, *J* = 5.3, 3.5 Hz, 1H), 3.87 (dd, J = 12.5, 2.6 Hz, 1H), 3.76 (d, J = 2.6 Hz, 1H), 3.72 (s, 3H). HRMS (ESI) m/z calcd for $C_{16}H_{18}N_3O_6S^+$ $[M+H]^+$ 380.0911, found 380.0913.

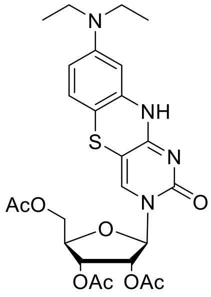

**2,3,5-tri-O-acetyl-$^{DEA}$tC-ribonucleoside.** The $^{DEA}$tC nucleobase (200 mg, 0.69 mmol) was suspended in anhydrous $CH_3CN$ (5 mL), and N,O-bis(trimethylsilyl)acetamide (254 µL, 1.04 mmol) was added. The reaction mixture was heated at 85°C under $N_2$ for 20 min and then allowed to cool to room temperature. 1,2,3,5-Tetra-O-acetyl-*β*-D-ribofuranose (265 mg, 0.83

mmol) and trimethylsilyl trifluoromethanesulfonate (151 μL, 0.83 mmol) were added. The reaction mixture was heated at 85°C for 2.5 h and allowed to cool to room temperature. The reaction progress was monitored with TLC (10% methanol in DCM). The reaction mixture was then extracted with 5% $NaHCO_3$ solution (25 mL) and $CH_2Cl_2$ (2 × 25 mL). After drying over anhydrous $Na_2SO_4$, the solvent was removed by rotary evaporation and the product was purified by flash chromatography on a Teledyne-ISCO CombiFlash Rf 200 (5% hexanes in EtOAc), yielding the semi-pure product as a yellow-orange oil that was carried through into deacetylation without further purification.

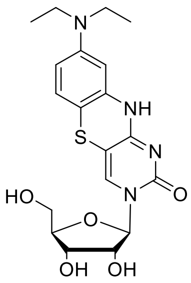

**$^{DEA}$tC-Ribonucleoside.** The semi-pure 2,3,5-Tri-O-acetyl-$^{DEA}$tC-ribonucleoside was dissolved in dry MeOH (4 mL) under $N_2$, and NaOMe (25 wt % in MeOH, 200 μL) was added. The reaction progress was monitored with TLC (10% methanol in DCM). After stirring for 1 h at room temperature, acetic acid (150 μL) was added, and the solvent removed by rotary evaporation. The crude product was dissolved in dichloromethane and purified by flash chromatography on a Teledyne-ISCO CombiFlash Rf 200 (20% MeOH in DCM), yielding the pure product as a yellow solid (136 mg, 64% over two steps). $^1$H NMR (500 MHz, DMSO-$d_6$) δ 7.90 (s, 1H), 6.80 (d, $J$ = 8.7 Hz, 1H), 6.43 (d, $J$ = 2.7 Hz, 1H), 6.28 (dd, $J$ = 8.7, 2.7 Hz, 1H), 5.73 (d, $J$ = 3.2 Hz, 1H), 5.31 (s, 1H), 5.15 (s, 1H), 4.97 (s, 1H), 3.95 (s, 2H), 3.83 (s, 1H), 3.69 (d, $J$ = 12.4 Hz, 1H), 3.56 (d, $J$ = 12.1 Hz, 1H), 3.25 (q, $J$ = 7.0 Hz, 4H), 1.05 (t, $J$ = 7.0 Hz, 6H). HRMS (ESI) m/z calcd for $C_{19}H_{25}N_4O_5S^+$ $[M+H]^+$ 421.1540, found 421.1543.

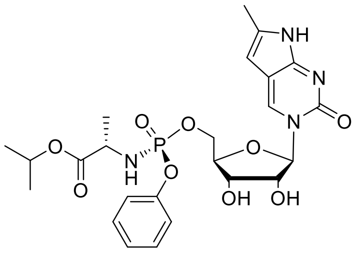

**Pyrrolo-C ProTide**. 1 M solution of tert-butylmagnesium chloride in THF (0.21 mL, 0.21 mmol) was added over a period of 3 minutes to a stirred suspension of Pyrrolo-C ribonucleoside (50 mg, 0.18 mmol) in dry THF (1 mL) at room temperature. The white suspension was stirred at this temperature for 30 min, and then (S)-Isopropyl 2-(((S)-(perfluorophenoxy)(phenoxy) phosphory)amino) propanoate (97 mg, 0.21 mmol) was added. The mixture was stirred at this temperature for 2 h. (S)-Isopropyl 2-(((S)-(perfluorophenoxy)(phenoxy)phosphory)amino) propanoate was added (97 mg, 0.21 mmol) and the reaction mixture was allowed to stir for 4 more hours. The reaction progress was monitored with TLC (5% methanol in DCM). The reaction mixture was extracted with dichloromethane (3 x 25 mL) in water (25 mL). After drying over anhydrous $Na_2SO_4$, the solvent was removed by rotary evaporation and the product was purified by flash chromatography on a Teledyne-ISCO CombiFlash Rf 200 (30% MeOH in DCM), yielding the product as a yellow solid (29 mg, 30%). There was no other isomer detectable by $^{31}P$ or $^{1}H$ NMR. $^{1}H$ NMR (400 MHz, DMSO-$d_6$) δ 11.09 (s, 1H), 8.25 (s, 1H), 7.37 (dd, $J$ = 8.4, 7.3 Hz, 2H), 7.27 – 7.14 (m, 3H), 6.07 (dd, $J$ = 13.0, 10.0 Hz, 1H), 5.95 (d, $J$ = 3.1 Hz, 1H), 5.76 (s, 1H), 5.56 (d, $J$ = 4.7 Hz, 1H), 5.21 (d, $J$ = 5.6 Hz, 1H), 4.83 (septet, $J$ = 6.2 Hz, 1H), 4.33 (m, 1H), 4.20 (m, 1H), 4.09 (m, 1H), 3.98 (m, 2H), 3.80 (m, 1H), 2.17 (s, 3H), 1.21 (d, $J$ = 7.1 Hz, 3H), 1.13 (d, $J$ = 6.3 Hz, 3 H), 1.12 (d, $J$ = 6.3 Hz, 3H. $^{31}P$ NMR (162 MHz, Chloroform-$d$) δ 7.22. $^{13}C$ NMR (101 MHz, Chloroform-$d$) δ 177.0, 162.4, 159.3, 154.2, 142.7, 137.6, 133.6, 129.0, 123.9, 114.9, 101.9, 96.8, 86.3, 79.3, 73.3, 72.7, 69.1, 54.2, 25.3, 25.2, 17.2. HRMS (ESI) m/z calcd for $C_{24}H_{32}N_4O_9P^+$ [M+H]$^+$ 551.1901, found 551.1903.

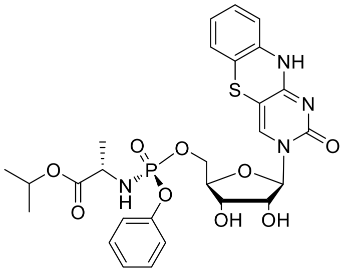

**tC ProTide**. 1 M solution of tert-butylmagnesium chloride in THF (0.34 mL, 0.34 mmol) was added over a period of 3 minutes to a stirred suspension of tC ribonucleoside (100 mg, 0.29 mmol) in dry THF (2 mL) at room temperature. The yellow suspension was stirred at room temperature for 30 min, and then (S)-Isopropyl 2-(((S)-(perfluorophenoxy)(phenoxy) phosphory)amino) propanoate (156 mg, 0.34 mmol) was added. The mixture was stirred for 4 h and monitored with TLC (5% methanol in DCM). The reaction mixture was extracted with dichloromethane (3 x 25 mL) in water (25 mL). After drying over anhydrous $Na_2SO_4$, the solvent was removed by rotary evaporation and the product was purified by flash chromatography on a Teledyne-ISCO CombiFlash Rf 200 (30% MeOH in DCM), yielding the product as a yellow solid (42 mg, 24%). There was no other isomer detectable by $^{31}$P or $^{1}$H NMR. $^{1}$H NMR (400 MHz, DMSO-$d_6$) δ 10.49 (s, 1H), 7.57 (s, 1H), 7.37 (t, $J$ = 7.8 Hz, 2H), 7.24 (d, $J$ = 8.0 Hz, 2H), 7.17 (t, $J$ = 7.4 Hz, 1H), 7.06 (t, J = 7.5, 1H), 7.01 – 6.87 (m, 3H), 6.02 (dd, $J$ = 13.2, 10.1 Hz, 1H), 5.75 (d, $J$ = 4.6 Hz, 1H), 5.44 (d, $J$ = 5.3 Hz, 1H), 5.22 (d, $J$ = 5.4 Hz, 1H), 4.85 (septet, $J$ = 6.3 Hz, 1H), 4.24 (m, 1H), 4.12 (m, 1H), 3.99 (m, 2H), 3.97 (m, 1H), 3.81 (m, 1H), 1.21 (d, $J$ = 7.1 Hz, 3H), 1.13 (d, $J$ = 6.3 Hz, 3H), 1.12 (d, $J$ = 6.3 Hz, 3H). $^{31}$P NMR (162 MHz, Chloroform-$d$) δ 3.39. $^{13}$C NMR (101 MHz, Chloroform-$d$) δ 173.6, 160.0, 155.7, 150.5, 135.4, 134.0, 129.6, 127.1, 125.7, 124.9, 124.4, 120.3, 117.6, 116.6,

97.7, 91.1, 82.8, 75.6, 69.8, 69.3, 65.6, 50.6, 21.6, 21.0. HRMS (ESI) m/z calcd for $C_{27}H_{32}N_4O_9PS^+$ $[M+H]^+$ 619.1622, found 619.1621.

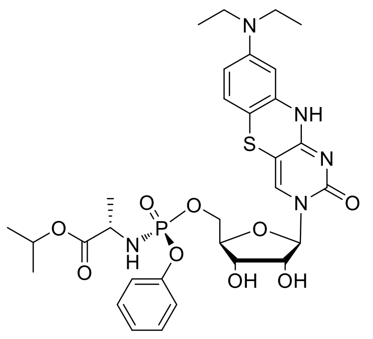

**$^{DEA}$tC ProTide**. 1 M solution of tert-butylmagnesium chloride in THF (0.4 mL, 0.4 mmol) was added over a period of 3 minutes to a stirred suspension of $^{DEA}$tC ribonucleoside (50 mg, 0.12 mmol) in dry THF (1 mL) at room temperature. The yellow suspension was stirred at room temperature for 30 min, and then (S)-Isopropyl 2-(((S)-(perfluorophenoxy)(phenoxy) phosphory)amino) propanoate (65 mg, 0.14 mmol) was added. The mixture was stirred for 4 h and monitored with TLC (5% methanol in DCM). The reaction mixture was extracted with dichloromethane (3 x 25 mL) in water (25 mL). After drying over anhydrous $Na_2SO_4$, the solvent was removed by rotary evaporation and the product was purified by flash chromatography on a Teledyne-ISCO CombiFlash Rf 200 (30% MeOH in DCM), yielding the product as a yellow solid (31 mg, 37%). There was no other isomer detectable by $^{31}$P or $^{1}$H NMR. $^{1}$H NMR (400 MHz, Chloroform-*d*) δ 9.07 (s, 1H), 7.59 (s, 1H), 7.25 (m, 5H), 7.09 (m, 1H), 6.54 (d, *J* = 8.7 Hz, 1H), 6.26 (d, *J* = 2.6 Hz, 1H), 6.20 (dd, *J* = 8.8, 2.5 Hz, 1H), 5.83 (d, *J* = 3.5 Hz, 1H), 4.95 (septet, *J* = 6.3 Hz, 1H), 4.88 (t, *J* = 10.8 Hz, 1H), 4.47 (m, 1H), 4.38 (m, 1H), 4.30 (m, 1H), 4.26 (t, *J* = 5.3 Hz, 1H), 4.10 (m, 2H), 3.24 (q, *J* = 7.1 Hz, 4H), 1.42 (d, *J* = 7.1 Hz, 3H), 1.20 (t, *J* = 6.3, 3H), 1.19 (t, *J* = 6.3, 3H),1.09 (t, *J* = 7.0 Hz, 6H). $^{31}$P NMR (162 MHz, Chloroform-*d*) δ 5.17. $^{13}$C NMR (101 MHz, Chloroform-*d*) δ 174.0, 160.7, 156.0, 150.7, 147.6, 136.5, 133.7, 129.7, 126.8, 125.0, 120.6, 108.5, 101.5, 100.6, 99.1, 91.6, 83.1,

75.9, 69.8, 69.4, 65.5, 50.9, 44.4, 21.8, 21.1, 12.7. HRMS (ESI) m/z calcd for $C_{31}H_{41}N_5O_9PS^+$ $[M+H]^+$ 690.2357, found 690.2356.

**Appendex**

$^1H$ NMR of Pyrrolo-C Ribonucleoside; 298K, DMSO, 400MHz:

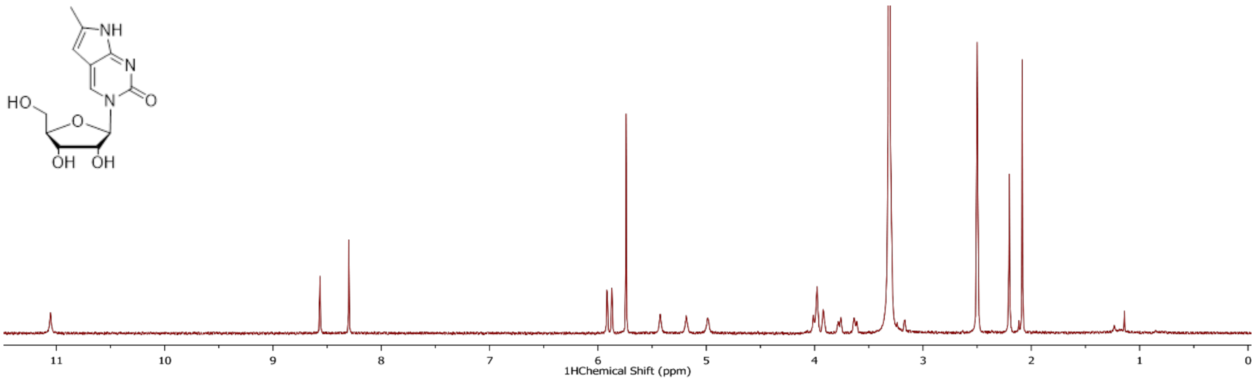


$^1H$ NMR of tC Ribonucleoside; 298K, DMSO, 400MHz

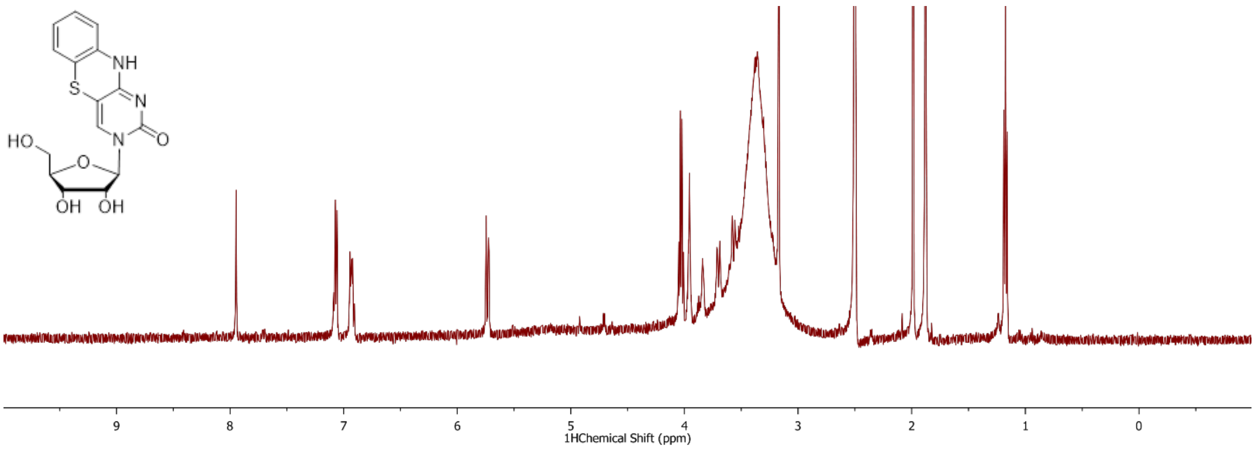

$^{1}H$ NMR of $^{MeO}$tC Ribonucleoside; 298K, $CD_3OD$, 400MHz

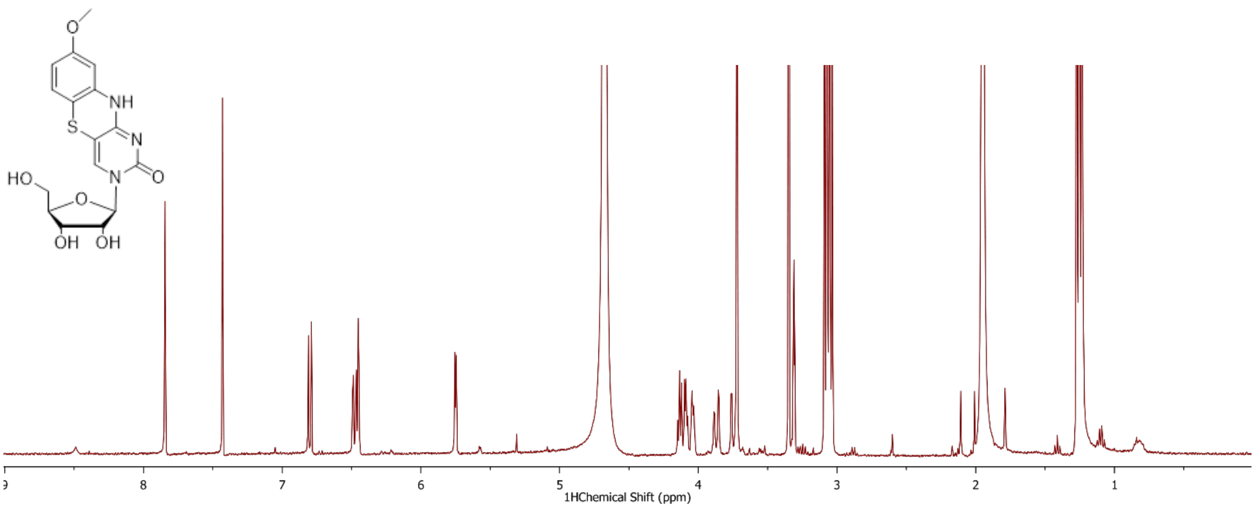


$^{1}H$ NMR of $^{DEA}$tC Ribonucleoside; 298K, DMSO, 400MHz

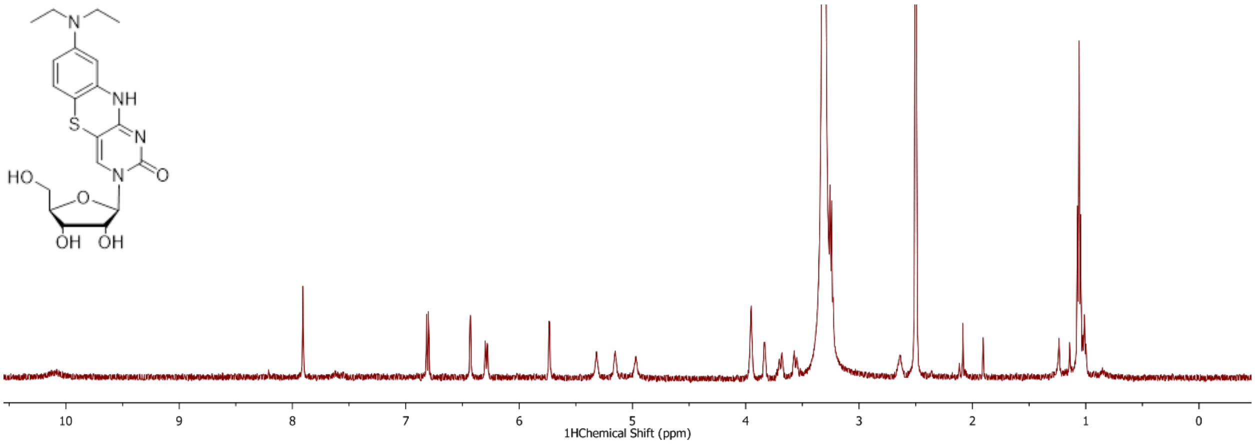

$^{1}H$ NMR, $^{13}C$ NMR and $^{31}P$ NMR of Pyrrolo-C ProTide; 298K, DMSO, 400MHz

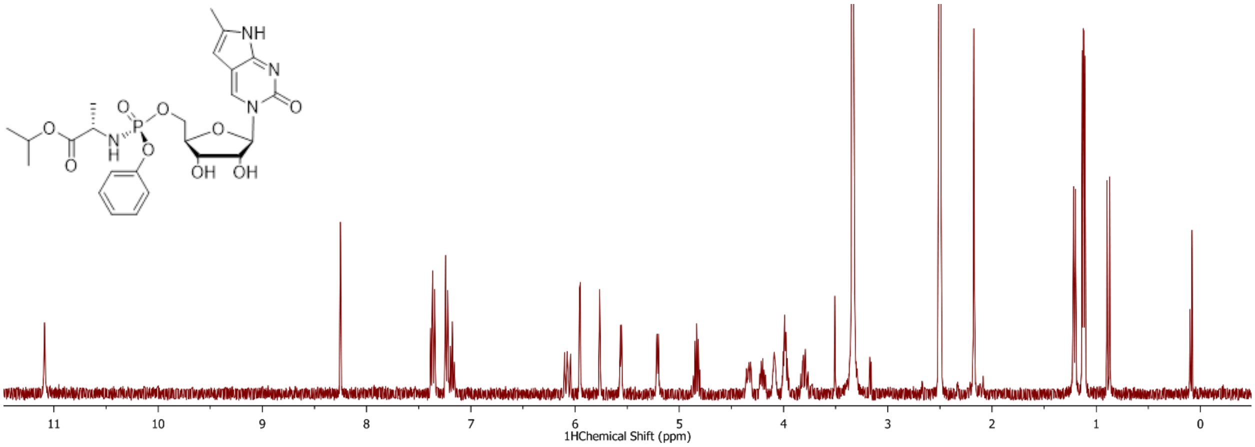


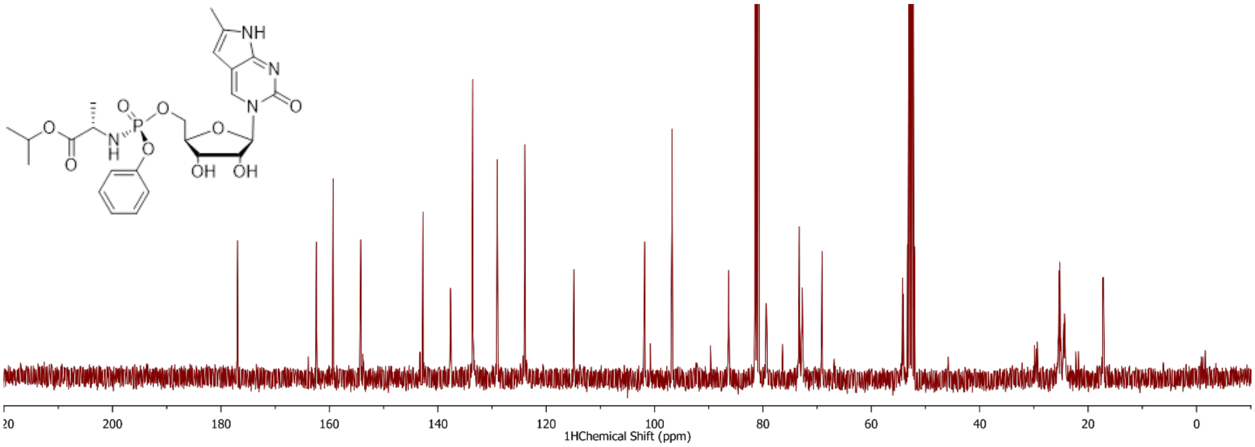


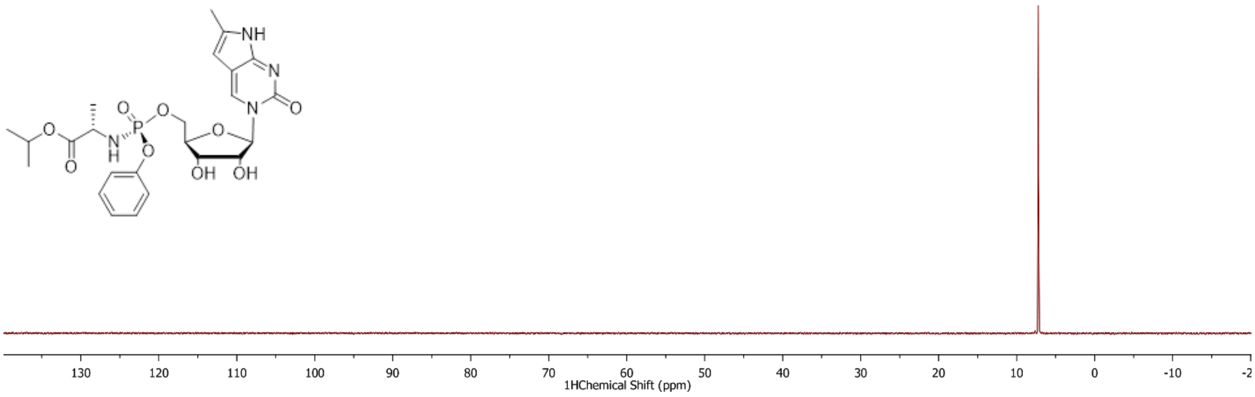

$^1$H NMR, $^{13}$C NMR and $^{31}$P NMR of tC ProTide; 298K, DMSO, 400MHz

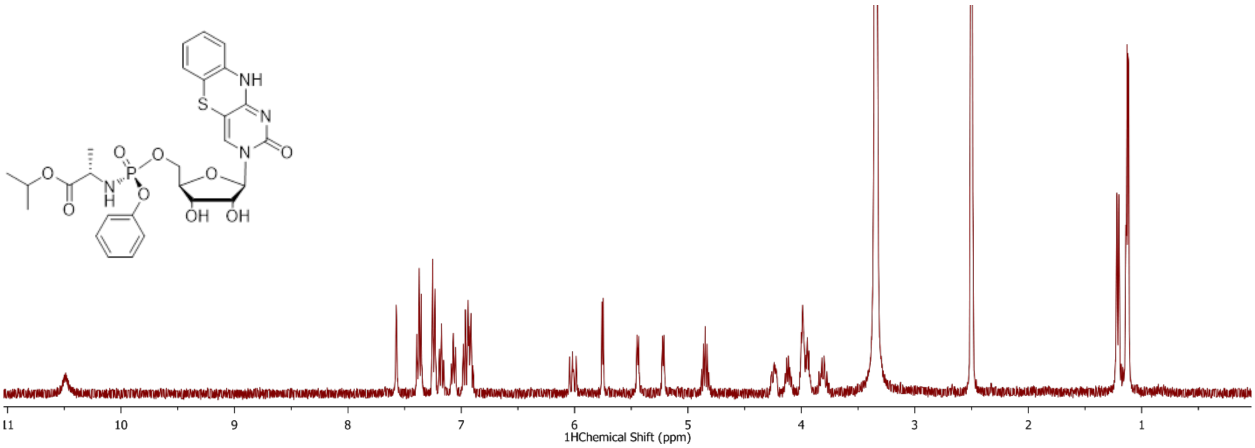


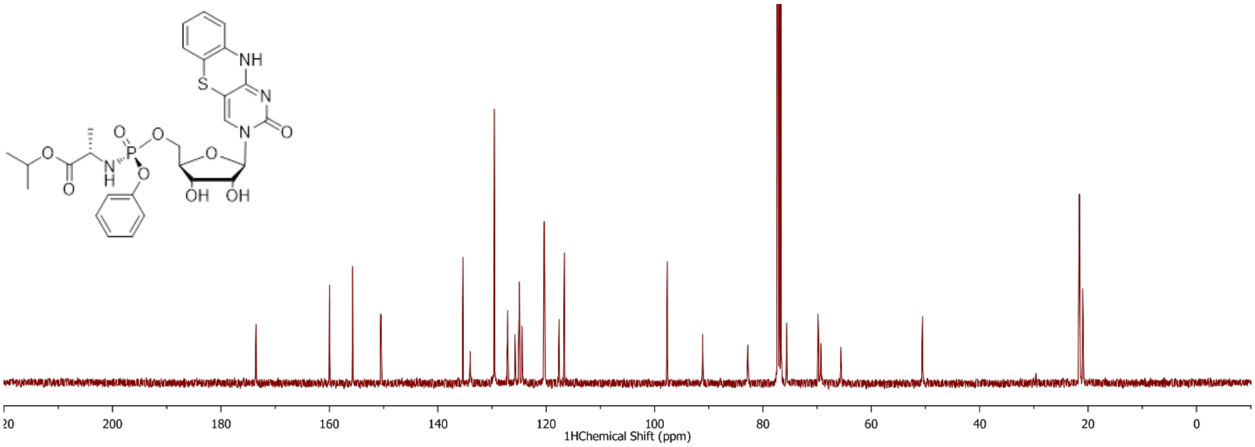


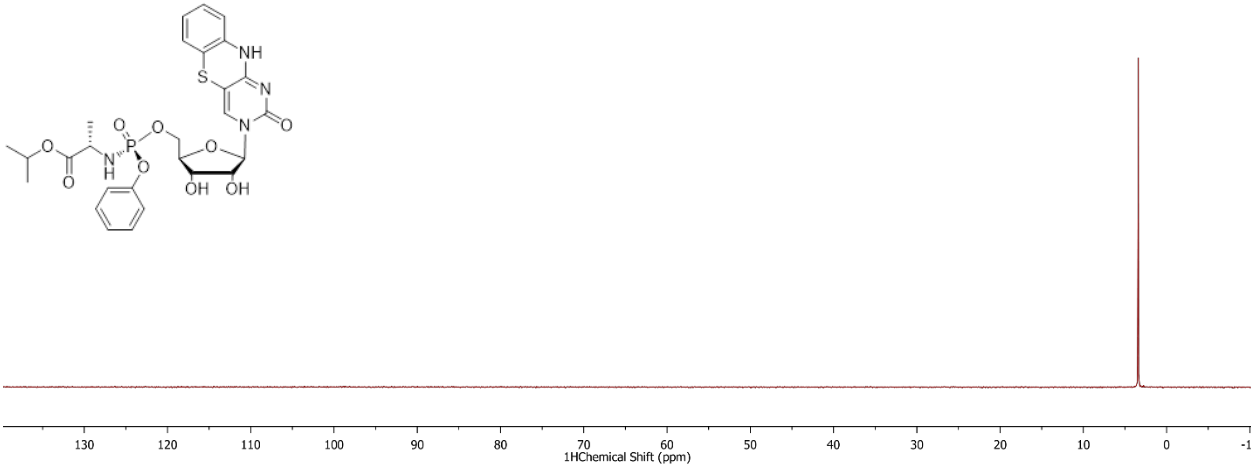

$^{1}$H NMR, $^{13}$C NMR and $^{31}$P NMR of $^{DEA}$tC ProTide; 298K, DMSO, 400MHz

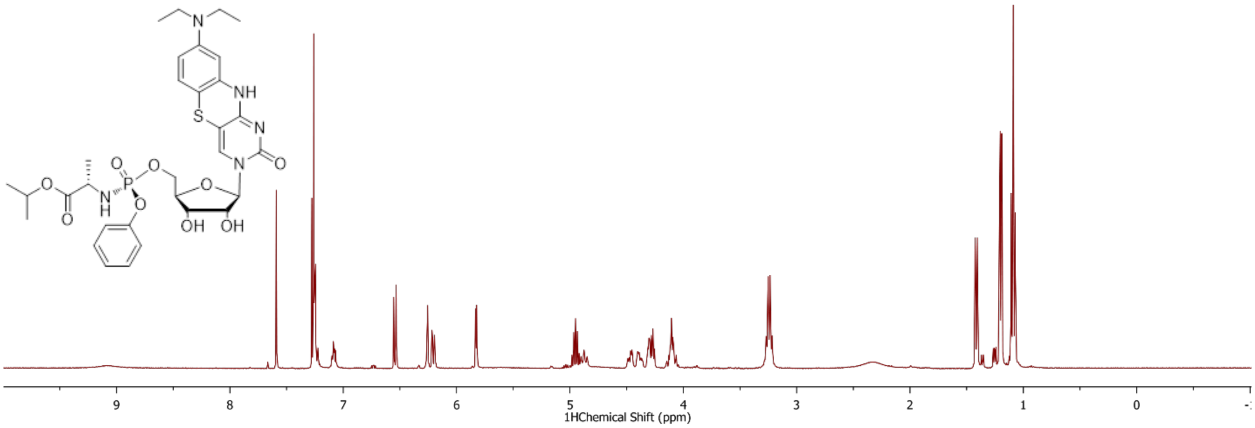


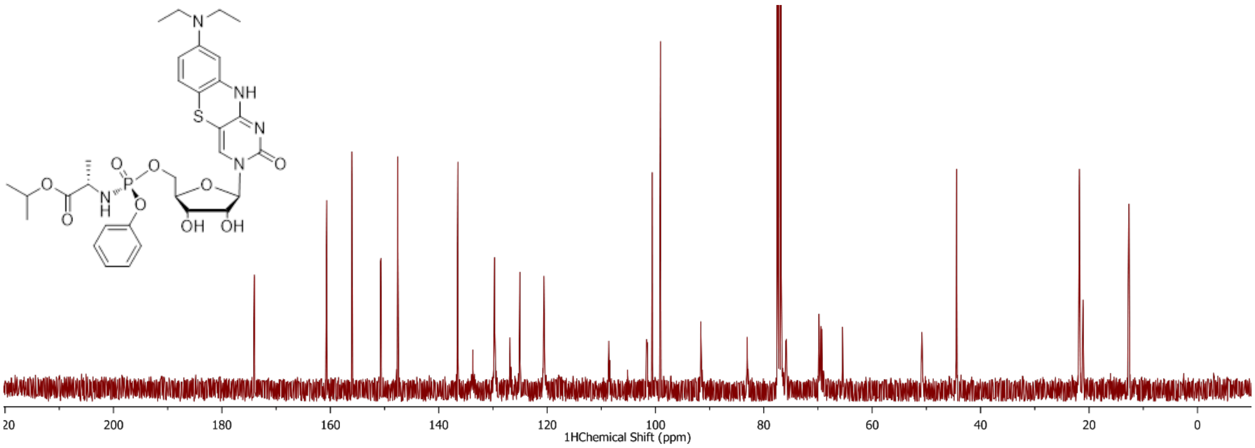


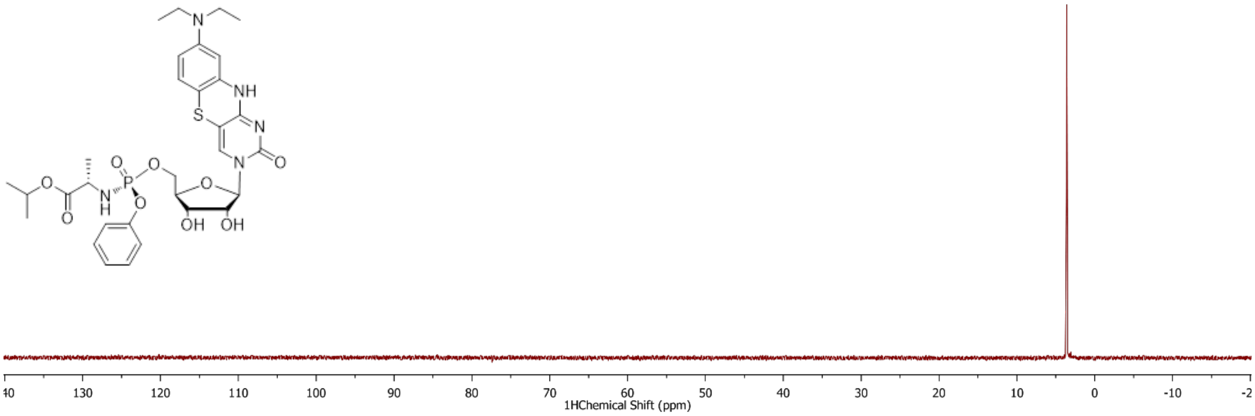

Quantum yield determination plots in 1× PBS buffer at 23 °C.

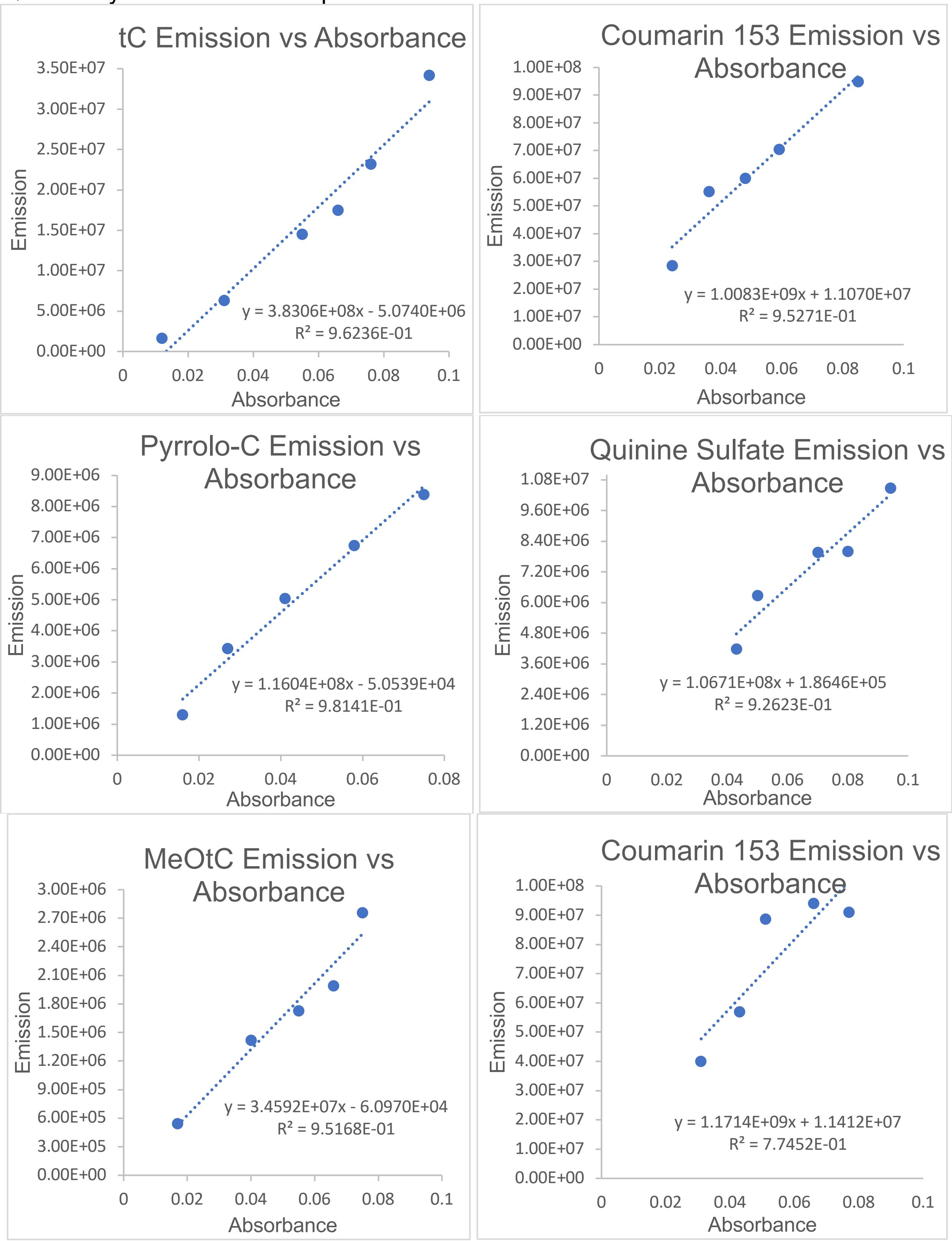

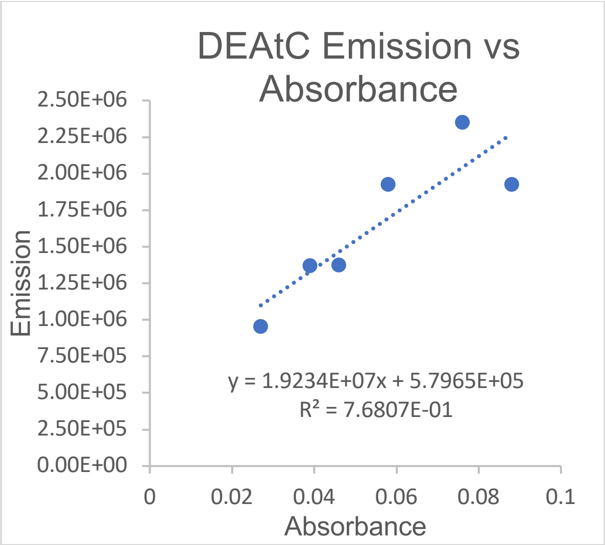
DEAtC Emission vs Absorbance
Emission
Absorbance
2.50E+06
2.25E+06
2.00E+06
1.75E+06
1.50E+06
1.25E+06
1.00E+06
7.50E+05
5.00E+05
2.50E+05
0.00E+00
0
0.02
0.04
0.06
0.08
0.1
y = 1.9234E+07x + 5.7965E+05
R² = 7.6807E-01

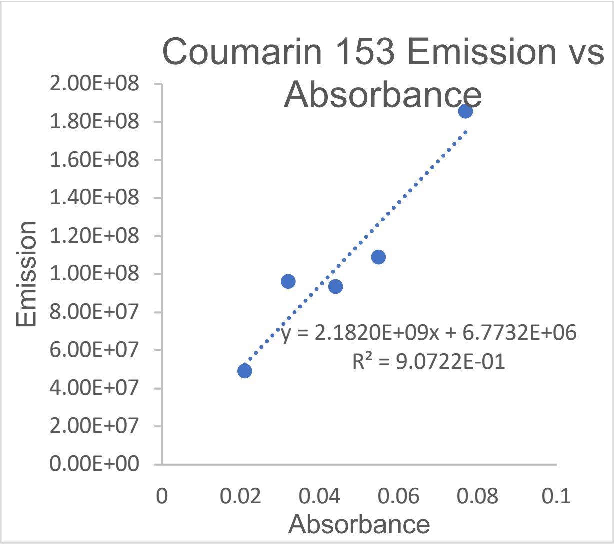
Coumarin 153 Emission vs Absorbance
Emission
Absorbance
2.00E+08
1.80E+08
1.60E+08
1.40E+08
1.20E+08
1.00E+08
8.00E+07
6.00E+07
4.00E+07
2.00E+07
0.00E+00
0
0.02
0.04
0.06
0.08
0.1
y = 2.1820E+09x + 6.7732E+06
R² = 9.0722E-01
